# Mechanobiological Specialization of Choroid Plexus Macrophages Defined by Titin Expression

**DOI:** 10.64898/2026.01.20.700716

**Authors:** Sagar Bhatta, Alayna Grzybowski, Gion Ortiz, Marcello DiStasio

**Affiliations:** Department of Ophthalmology and Visual Science, Yale School of Medicine, New Haven, CT; Department of Pathology Yale School of Medicine, New Haven, CT; Program in Forensic Science, University of New Haven, New Haven, CT

**Author notes:** Correspondence to Marcello DiStasio, 300 George Street, New Haven, CT, 06511.

**Keywords:** Choroid plexus, Border-associated macrophages, Titin (TTN), Cytoskeletal remodeling, Mechanotransduction, Alzheimer’s disease, Single-nucleus RNA sequencing, Spatial transcriptomics

## Abstract

Functioning both as the site of cerebrospinal fluid production and as an interface between the peripheral circulation and the central nervous system, the choroid plexus (ChP) modulates the gating of immune cells and signals from the periphery as they pass from the blood into cerebrospinal fluid. We generated a single-nucleus and spatial transcriptomic atlas of the adult human ChP, in donors with and without Alzheimer’s disease (AD), revealing transcriptionally and spatially distinct macrophage states. We identified a previously unrecognized population of titin-expressing (TTN^+^) tissue-resident macrophages characterized by coordinated activation of cytoskeletal remodeling, phagocytosis, autophagy, and inflammatory programs. Isoform-specific qPCR, protein immunofluorescence, and single-molecule FISH validated TTN expression in situ. In AD, TTN^+^ macrophages expand and undergo broad transcriptional rewiring, including activation of MEF2-linked mechanotransduction pathways, elevated senescence signatures, and a loss of ligand–receptor connectivity with epithelial, endothelial, and stromal partners. Collectively, our findings define a mechanosensitive macrophage program at the blood–CSF interface and uncover a TTN-associated shift in macrophage function and microenvironment integration in AD.

## 1 Introduction

The choroid plexus (ChP) is a vital brain structure consisting of a network of capillaries surrounded by mesenchymal and epithelial cells that reside within the brain’s ventricles. The ChP produces cerebrospinal fluid (CSF) and, along with arachnoid membranes, forms the blood-CSF barrier. This specialized barrier and signaling hub exists at the interface between the peripheral circulation and the central nervous system (CNS). The ChP functions as an active immunological checkpoint that integrates peripheral signals with CNS-derived cues to regulate leukocyte entry, antigen presentation, and inflammatory tone^1^. Anatomically and functionally, the ChP resides at a strategic neuroimmune nexus: resident stromal, epithelial, and endothelial cells form a scaffold for diverse immune populations that continuously survey the ventricular environment and coordinate immune responses across the blood–CSF interface.

Among these immune populations, ChP macrophages exhibit a hybrid identity. They share features with CNS border-associated macrophages (BAMs), circulating monocytes, and tissue-resident macrophages, and display functional diversity shaped by stromal niche cues, barrier architecture, and CSF-derived cytokines^2,3,4^. Despite their central role in neuroimmune communication, the full molecular diversity of ChP macrophages in the adult human brain has not been defined, and the mechanisms that govern their differentiation and functional states remain poorly understood.

A significant gap in understanding concerns how ChP macrophages integrate mechanical signals, a dimension of immune regulation increasingly recognized across tissues^5^. Mechanical forces shape leukocyte migration, adhesion, antigen scanning, and inflammatory activation; cytoskeletal programs are central to how immune cells sense and respond to physical cues^6,7,8^. Macrophage polarization is known to be influenced by biophysical cues in the extracellular environment such as interstitial flow, stretching, spatial confinement, and matrix stiffness^9^. Our discovery of titin (TTN) expression in human ChP macrophages reveals an unexpected component of the mechanobiological axis in a CNS border compartment.

TTN is the largest protein in the human proteome and a canonical organizer of sarcomere elasticity in muscle^10^, but recent work demonstrates that TTN also contributes to immune cell mechanics: human T cells express multiple TTN isoforms, and TTN regulates microvillar architecture, integrin activation, and chemotactic efficiency^11^. TTN-deficient lymphocytes display impaired motility and altered actin dynamics, identifying TTN as a master regulator of immune cell trafficking. These observations suggest that TTN operates as a mechanotransductive scaffold that integrates cytoskeletal tension, cell–environment interactions, and signal responsiveness, processes central to macrophage behavior within mechanically heterogeneous tissues like the ChP.

Mechanistically, TTN’s modular domains can buffer mechanical strain, bind the actin–myosin cytoskeleton, and interact with regulatory complexes that engage MEF2 transcription factors, which themselves serve as mechanoresponsive sensors downstream of cytoskeletal deformation and calcium signaling^12,5^. Consistent with this, our data show that TTN^+^ ChP macrophages display elevated MEF2A/C/D activity and upregulation of pathways involved in cytoskeletal remodeling, phagocytosis, oxidative stress responses, MHC class II antigen presentation, and autophagy. These signatures collectively point towards a macrophage phenotype optimized for tension-bearing, motility, barrier engagement, and dynamic interactions with stromal and epithelial folds. In the densely folded, fluid-exposed, and mechanically active environment of the ChP, TTN expression may therefore provide a structural and regulatory advantage for macrophages navigating stromal surfaces, engaging epithelial barriers, and responding to blood and CSF flow–derived shear forces.

Mechanobiological signaling may play a key role in the immunobiology of Alzheimer’s disease (AD), a condition increasingly conceptualized as a disorder of neuroimmune imbalance and barrier dysfunction^13^. While microglial activation in the parenchyma is well established^14,15^, the ChP represents an understudied immunological interface that undergoes structural, transcriptional, and inflammatory remodeling in aging and AD^16,17^. ChP epithelial cells, stromal fibroblasts, and macrophages work together to regulate leukocyte recruitment and immune priming. Disruptions in these processes may alter neuroimmune signaling, impair CSF-mediated waste clearance, and contribute to chronic inflammation^18^. Our findings demonstrate that TTN^+^ macrophages are significantly expanded in AD tissue and adopt a senescent, spatially segregated, and interaction-deficient state, with reduced ligand–receptor connectivity to epithelial, endothelial, and stromal partners.

These observations position TTN as a previously unrecognized regulator of macrophage mechanobiology at the brain’s borders. They raise the possibility that TTN-dependent cytoskeletal programs shape macrophage behavior within the ChP, and therefore, disruptions to this axis contribute to impaired immune surveillance, altered barrier signaling, and chronic inflammation in Alzheimer’s disease.

## 2 Results

### A single-nucleus transcriptomic atlas defines the cellular landscape of the human choroid plexus

To characterize the cellular architecture of the adult human choroid plexus (ChP), we generated a single-nucleus RNA-sequencing (snRNA-seq) atlas from postmortem lateral ventricular ChP tissue from donors with (n=4) and without (n=5) Alzheimer’s disease (AD). Integration of data across donors and technical variability was mitigated using scVI^19^ for batch correction and latent space integration, enabling robust alignment of transcriptomic signatures. Dimensionality reduction and visualization were performed with scanpy^20^, using the integrated scVI latent space to generate high-resolution UMAP embeddings that revealed distinct macrophage subclusters and their transcriptomic heterogeneity. Cell-type annotation was performed using a custom, manually curated panel of ChP–specific marker genes (see Supplementary Table S2), allowing accurate delineation of epithelial, stromal, neuronal, and immune populations.

UMAP projection of expression profiles from 37,378 nuclei revealed major epithelial, endothelial, mesenchymal, immune, glial, and neuronal compartments (Figure 1A). Bayesian compositional modeling using scCODA^21^ demonstrated that epithelial cells dominate the tissue (59.5%), followed by endothelial and mesenchymal populations, with immune lineages including macrophages, dendritic cells, B cells, and T cells, comprising ~14% of total nuclei (Figure 1B). Notably, the diagnosis of AD did not significantly alter the baseline proportional representation of any cell class, as reflected by insignificant coefficients in the scCODA model.

**Figure 1.**
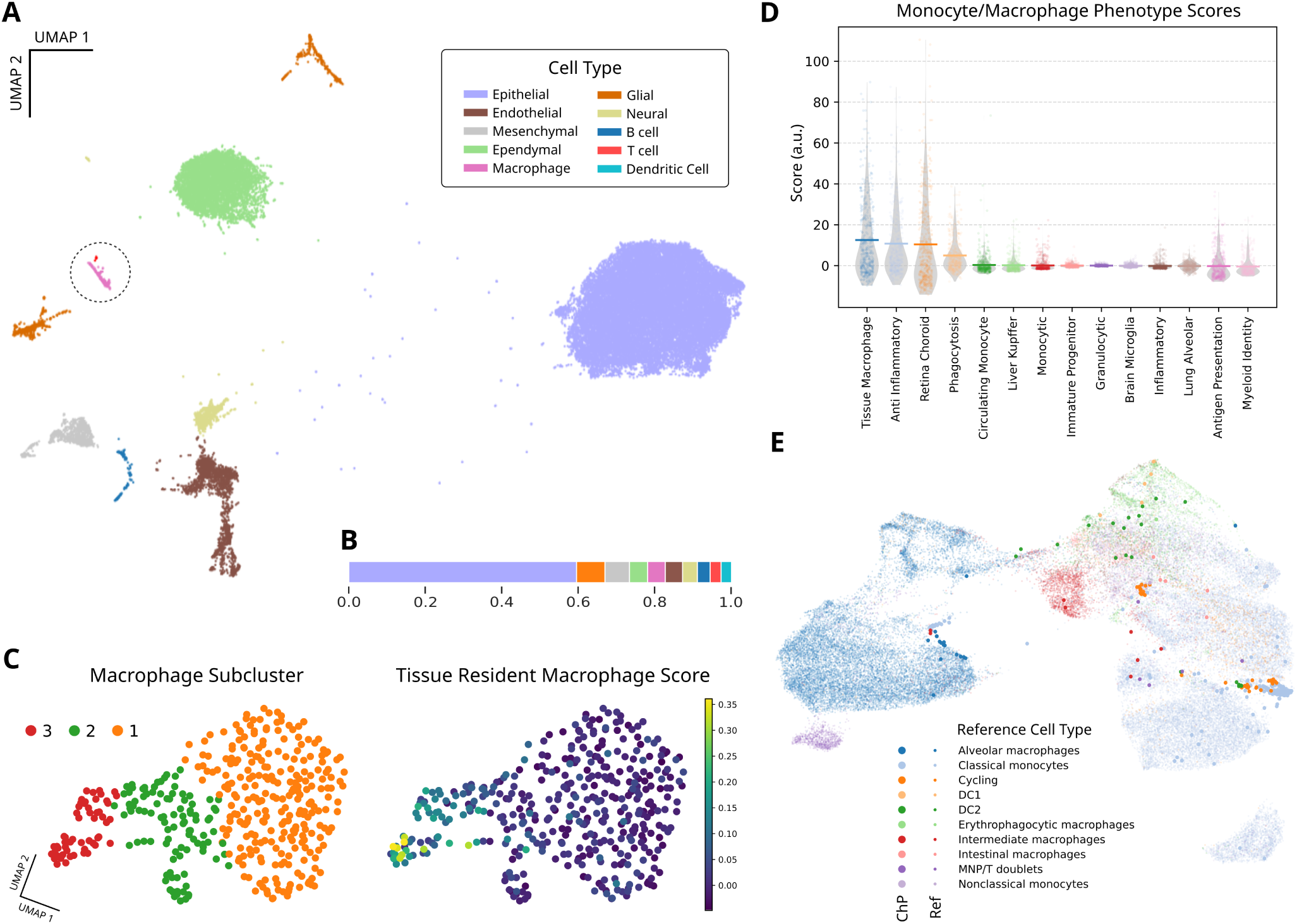
Cell type composition of human choroid plexus. **(A)** Uniform Manifold Approximation and Projection (UMAP) of all single-nucleus transcriptomes profiled from ChP tissue. Distinct clusters correspond to major choroid plexus compartments, including epithelial cells, endothelial and vascular-associated cells, mesenchymal/stromal populations, immune lineages (macrophages, dendritic cells, lymphocytes), glial and ependymal-like cells, and rare neuronal nuclei. Colors indicate curated cell type assignments based on lineage markers and differential expression analyses. **(B)** Inferred underlying proportions of each cell class, distribution, derived from intercepts of an scCODA model fit to the complete dataset accounting for compositional constraints and batch effects in the data, indicating that epithelial cells dominate the tissue com-position (59.5%), followed by endothelial (7.4%), mesenchymal (6.4%), macrophage (4.6%), glial (4.5%), ependymal (4.7%), neural (3.9%), B-cells (3.3%), dendritic cells (2.7%), and T-cells (2.9%). **(C)** UMAP visualization of macrophage subclusters and tissue-resident macrophage scores. Left: Single-cell UMAP projection of macrophages, colored by manually annotated subclusters (Cluster 1: orange; Cluster 2: green; Cluster 3: red), highlighting transcriptionally distinct populations. Right: Same UMAP projection showing the tissue-resident macrophage (TRM) score per cell, with color intensity corresponding to TRM signature enrichment (scale from 0 [low, purple] to 0.35 [high, yellow]), indicating enrichment of TRM features in specific subclusters. **(D)** Monocyte/Macrophage Phenotype Scores. Violin plots show enrichment scores for curated myeloid gene sets representing monocyte and macrophage phenotypes across diverse tissues. Individual points indicate sample-level scores, with horizontal bars marking median values. **(E)** ChP macrophages mapped onto Cross-Tissue Immune Atlas myeloid compartment. Each dot represents an individual cell, with colors indicating different cell types; larger dots represent ChP macrophages in our dataset.

### Choroid plexus macrophages are a heterogeneous population with characteristics of border-associated macrophages

Subclustering of macrophages (n=412) resolved three transcriptionally distinct groups, each exhibiting differential enrichment for tissue-resident macrophage (TRM) signatures (Figure 1C). To contextualize these states, we used scanpy^22^ to compute module scores spanning diverse myeloid programs—including TRM identity, inflammatory activation, phagocytosis, metabolic states, and antigen presentation—which revealed substantial functional heterogeneity across clusters (Figure 1D; for a listing of genes used to define these modules, see Supplementary Table S3). Mapping these cells onto a cross-tissue myeloid atlas^23^ showed that ChP macrophages occupy transcriptional positions bridging classical monocytes, intermediate macrophages, alveolar-like macrophages, and dendritic cell states (Figure 1E), consistent with a composite peripheral-CNS border identity. The majority of the macrophages from the ChP dataset mapped to classical monocytes and cycling cells. Cells with signatures similar to alveolar macrophages, DC2 (dendritic cells), and intermediate macrophages also represented significant portions of the mapped data. Additionally, smaller subsets of cells mapped to intestinal macrophages and erythrophagocytic macrophages, highlighting the presence of gene signatures that resemble those found in these specialized macrophage types. This mapping suggests the existence of a variety of macrophage subtypes within the ChP that express gene signatures similar to immune cells across different tissues.

Differential expression across macrophage subclusters reveals gene sets that segregate into three transcriptionally coherent modules reflecting distinct functional identities within the choroid plexus (Figure 2A). Subcluster 1 contains signatures of barrier associated functions such as epithelial-interaction and homeostatic tissue-residence (e.g. CX3CR1, SPIC, MUC16, CLSTN2, CASR, HYKK). Subcluster 2 comprises macrophages enriched for tissue-resident surveillance and immune quiescence (e.g. CX3CR1, SPP1, and TNFRSF14), metabolic and mitochondrial specialization (e.g. MRPL17, ECHS1, and MT1H), and proteostatic and chromatin regulatory programs (e.g. UBQLN2, HMGN2, SAP25), suggestive of stress-adaptive or remodeling functions consistent with long-lived, interface-associated macrophages. Subcluster 3 is enriched for genes involved in anti-inflammatory and pro-resolving activity (e.g. ANXA1, SUMO4), tissue remodeling and matrix interaction (e.g. LYVE1, SPP1), RNA-processing and stress adaptation (e.g. PRPF4, HNRNPUL2), and vesicular and ion-regulatory dynamics (STX16, CLIC2). Human ChP macrophages thus show divergent structural, metabolic, and immune states similar to those in mice^24^ and other sites of tissue residence.

**Figure 2.**
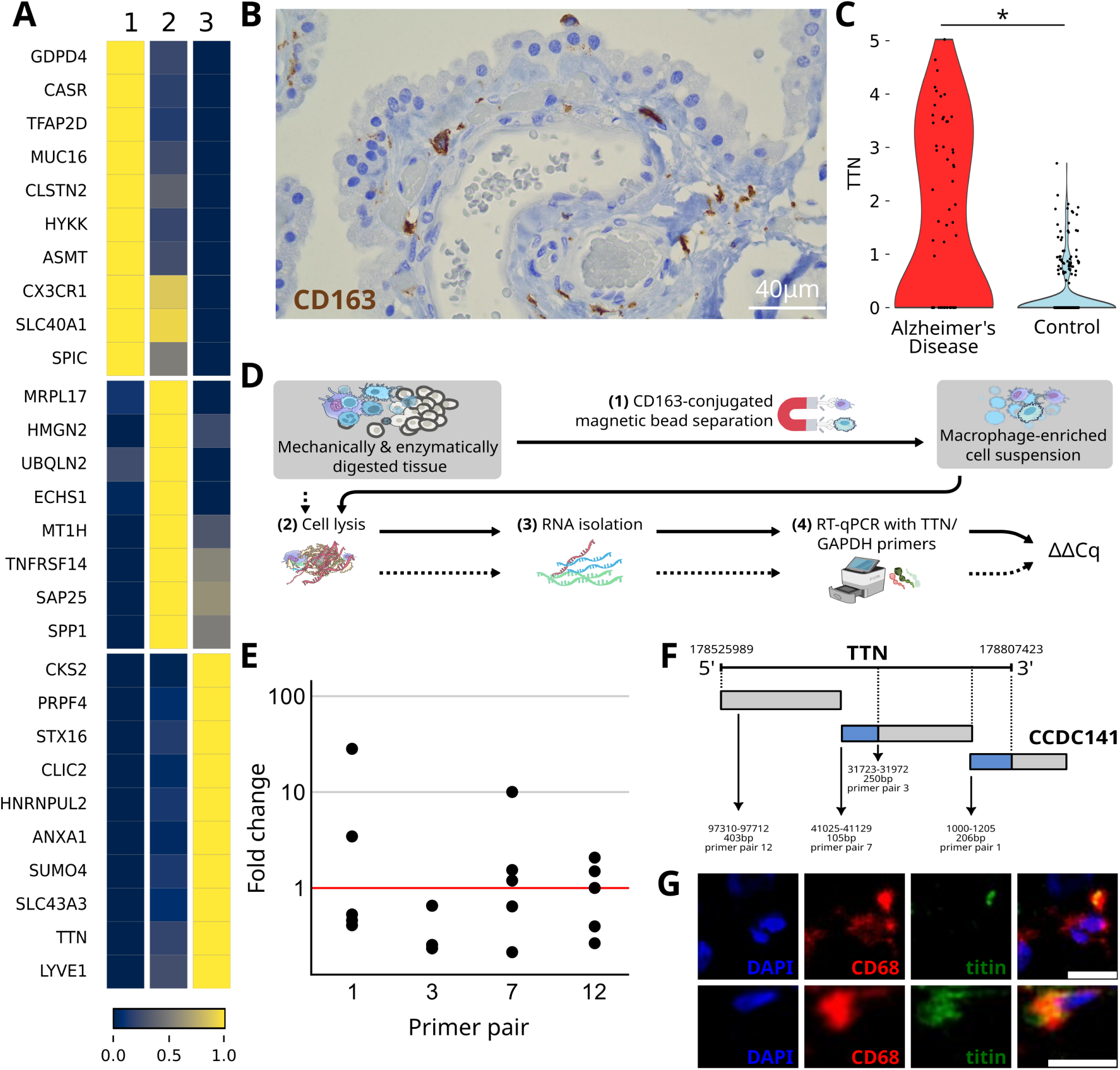
Titin-expressing macrophages are present in the choroid plexus. **(A)** Heatmap of the top eight differentially expressed genes in each of the three ChP macrophage sub-clusters. Gene expression levels are represented on a normalized gradient scale. **(B)** DAB immunohistochemical stain of a 5 µm FFPE section of control ChP tissue using a CD163 antibody. CD163-positive macrophages are marked by brown staining, while cell nuclei are counterstained blue with hematoxylin, indicating their distribution within the vascularized ChP tissue. Scale bar = 40 µm. **(C)** Violin plots illustrating TTN expression levels in ChP samples from Alzheimer’s disease and control conditions. Each plot shows the distribution, density, and variability of TTN transcript counts in ChP macrophages within each condition. **(D)** Schematic representation of the experimental workflow for evaluation of TTN RNA enrichment in macrophages, involving tissue dissociation, CD163 positive cell enrichment, RNA isolation, cDNA synthesis, PCR amplification using isoform-specific TTN primers, and ΔΔCq calculation with GAPDH used for normalization. **(E)** Fold changes, computed from ΔΔCq, in TTN isoform expression relative to GAPDH in CD163-bead enriched macrophages relative to whole tissue. The x-axis represents different primer sets (1, 3, 7, and 12), and the y-axis shows fold changes (log scale). Each point represents a single donor sample/primer set pair. **(F)** Schematic representation of the TTN gene with highlighted primer binding sites, with genomic coordinates on chromosome 2, indicating selected splicing sites and isoforms of TTN. **(G)** Immunofluorescence images of ChP with anti-CD68 (red) and anti-titin (green) primary antibodies, co-stained with DAPI (blue), showing cytoplasmic titin protein in ChP macrophage cytoplasm. Merged images are in right column. Scale bars = 20 µm.

### Choroid plexus macrophages include a titin-expressing population validated by isoform-specific qPCR

TTN emerged as a distinguishing transcript of ChP macrophage subcluster 3 (Figure 2A). TTN^+^ macrophages constituted 21.8% of the total macrophage population. Immunohistochemistry confirmed the presence of CD163^+^ macrophages embedded within the stromal and epithelial folds of the ChP (Figure 2B). Violin plots of distributions of TTN expression in single macrophages demonstrated significantly higher levels in AD compared with control samples (Figure 2C). Expression of TTN RNA was evaluated in publicly available single-cell RNA-sequencing atlases, identifying TTN^+^ myeloid cells in multiple datasets (see Supplementary Figure S1A and B). TTN expression was particularly prominent in myeloid/macrophage cells occupying adipose tissue (see Supplementary Figure S1C).

To validate TTN expression independently of sequencing, we isolated CD163^+^ macrophages and performed isoform-specific RT-qPCR for four regions of the TTN transcript. Primer sets 1, 3, 7, and 12 detected TTN transcripts at variable abundance in CD163^+^ cells, with primer pairs targeting the N-terminal and C-terminal regions yielding robust amplification (Figure 2D–F) relative to a GAPDH control primer set. These results support the existence of TTN-expressing macrophages in human ChP tissue and suggest isoform diversity consistent with cell-type–specific splicing. TTN expression was also confirmed on RT-qPCR of RNA extracted from macrophages isolated from frontal cortex, CSF, subcutaneous fat, and mesenteric fat (see Supplementary Figure S2). Additionally, immunofluorescence staining of post-mortem human ChP samples demonstrated titin protein in the cytoplasm of CD68^+^ macrophages (Figure 2G).

### TTN^+^ macrophages exhibit distinct molecular programs and occupy altered spatial niches in the choroid plexus

We next asked whether TTN expression defines a functionally distinct macrophage state. Gene signature scoring using scanpy^22^ revealed that TTN^+^ cells display coordinated upregulation of cytoskeletal remodeling, phagocytic, lysosomal, autophagy, inflammatory, and oxidative stress-response pathways, as well as elevated DAM-like and senescence signatures (Figure 3A; gene lists for modules are given in Supplementary Table S4). Given the close link between TTN and mechanotransduction, we examined expression of MEF2A, MEF2C, and MEF2D, transcription factors with known roles in cytoskeletal regulation and stress adaptation. All three showed enrichment in the TTN^+^ macrophage subcluster (Figure 3B), supporting a possible TTN–MEF2 axis in ChP macrophages. These patterns suggest that TTN^+^ macrophages occupy a mechanically and metabolically engaged phenotype.

**Figure 3.**
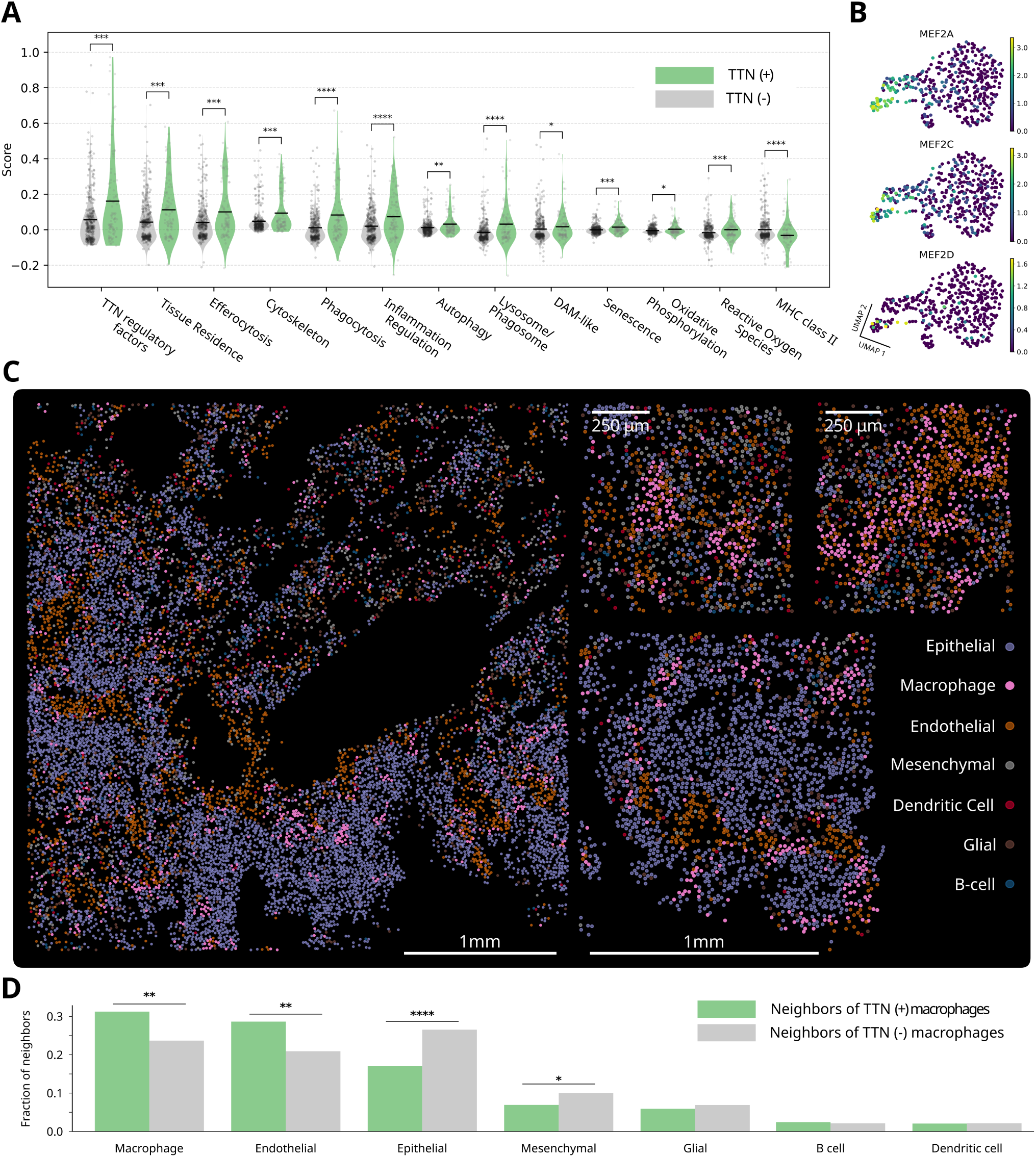
Spatial organization and functional states of TTN-expressing ChP macrophages **(A)** Distinct molecular programs are expressed in TTN^+^ macrophages. Violin plots showing module scores for curated macrophage gene signatures. TTN^+^ cells exhibit enhanced cytoskeletal, phagocytic, autophagy-lysosomal, inflammatory, oxidative stress–response, and MHC class II programs, along with elevated senescence and DAM-like signatures, compared with TTN-macrophages. **(B)** MEF2 transcription factor activity in macrophage subclusters. Spatially resolved expression of MEF2A, MEF2C, and MEF2D highlights differential activation across macrophage populations, consistent with transcriptional profiles associated with cytoskeletal remodeling and mechanosensitive signaling. **(C)** Spatial distribution of immune and stromal populations in the choroid plexus. High-resolution spatial transcriptomics maps show the organization of epithelial, endothelial, mesenchymal, macrophage, dendritic cell, glial, and B-cell compartments. TTN^+^ macrophages localize within regions of dense epithelial–immune interface. Scale bars, 1 mm and 250 µm. **(D)** Altered local cellular neighborhoods of TTN^+^ macrophages. Bar plots quantifying the proportion of neighboring cell types surrounding TTN^+^ versus TTN-macrophages demonstrate that TTN^+^ cells engage distinct spatial niches, with increased macrophage-endothelial and macrophage–macrophage proximity and reduced interactions with epithelial and mesenchymal compartments.

Spatial transcriptomics using slide-seq^25^ on ChP from 4 donors with AD, with integration of our snRNA-seq data via GIMVI^19^, revealed extensive local vascular-immune and epithelial–immune interaction within the ChP folds and demonstrated that TTN^+^ macrophages occupy distinct microenvironments enriched for endothelial and immune interfaces (Figure 3C). Neighborhood analysis confirmed that TTN^+^ macrophages preferentially cluster near other macrophages and show reduced proximity to epithelial and mesenchymal cells relative to TTN-cells (Figure 3D). These spatial shifts indicate altered tissue integration and microenvironmental sensing associated with TTN expression.

### Alzheimer’s disease drives transcriptional remodeling, senescence, and loss of interaction networks in choroid plexus macrophages

To define AD-associated remodeling, we performed differential expression analysis between AD (n=63) and control (n=349) macrophages. AD macrophages exhibited a broad transcriptional shift, with TTN, STAB1, SAT1, MEF2A, and TNFAIP3 among the most significantly upregulated genes, while neuronal-adhesion and cytoskeletal-associated transcripts (RBFOX1, PTPRD, DLG2, CSMD1) were prominently downregulated (Figure 4A). These changes highlight disrupted cytoskeletal, stress-response, and immune-modulatory programs in AD macrophages.

**Figure 4.**
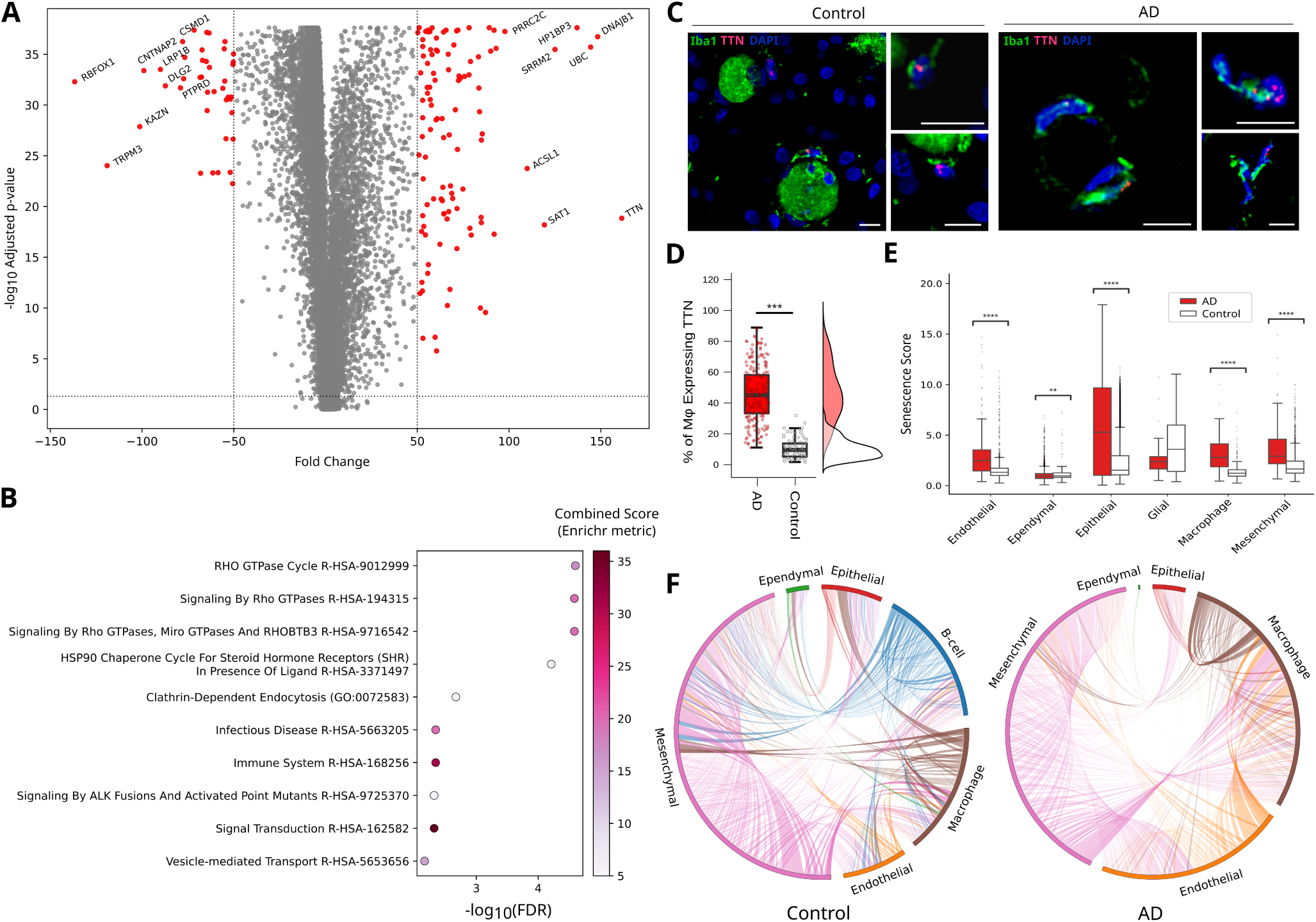
Alzheimer’s disease reshapes transcriptional state, senescence programs, and interaction networks of choroid plexus macrophages. **(A)** Global transcriptional changes in choroid plexus macrophages in Alzheimer’s disease. Volcano plot depicting differentially expressed genes (DEGs) between AD and control choroid plexus macrophages. For each gene, the fold change (x-axis) is plotted against the -log_10_ adjusted p-value (y-axis). Genes surpassing the significance thresholds (adjusted p-value *<*0.05 and |FC*| >*50) are highlighted in red, while dashed lines denote the statistical and fold-change cutoffs. Representative genes with the largest effect sizes or strongest statistical significance are annotated. **(B)** Over-representation analysis was performed using Enrichr across GO Biological Process (2023), KEGG (2021), and Reactome (2022) gene sets. The dot plot summarizes the top enriched pathways, with the x-axis representing enrichment significance (–log_10_ adjusted p value) and dot color reflecting the Enrichr combined score, which integrates both statistical significance and deviation from background expectation. Pathways are ranked by adjusted P value, highlighting the major biological processes associated with the observed gene expression changes. **(C)** Example micrographs of macrophages in control and Alzheimer’s disease choroid plexus, stained with Iba1 immunofluorescence (green), RNAscope single molecule FISH probes for TTN RNA (red), and DAPI nuclear stain (blue). All scale bars = 10 µm. **(D)** Frequency of macrophages containing TTN transcripts in RNAscope assay, quantified as percentage of total number of macrophages present per field. An increased fraction of TTN-expressing macrophages are found in AD ChP compared to control (*** = p *<*0.001, Bayesian independent samples t-test reporting posterior probability). **(E)** Senescence scores derived using the senepy framework demonstrate increased senescence signatures across multiple choroid plexus cell types, with the most pronounced elevation in macrophages. **(F)** Circos plots of ligand–receptor interactions inferred by LIANA show reduced macrophage connectivity to epithelial, endothelial, and stromal populations in AD, indicating impaired integration into the choroid plexus niche.

Pathway enrichment analysis (Figure 4B) strongly implicated dysregulation of Rho family GTPase signaling, a core regulatory axis that integrates mechanical inputs with cytoskeletal remodeling, adhesion dynamics, and migratory behavior. RhoA, Rac1, and Cdc42 act as molecular switches that coordinate actin polymerization, membrane protrusion, and contractility—programs essential for macrophage motility, phagocytosis, and environmental sensing, consistent with altered mechanotransduction and responsiveness to tissue cues. Additional perturbations are seen in clathrin-dependent endocytosis, HSP-90 chaperone activity, vesicle trafficking, and immune activation pathways, consistent with impaired environmental responsiveness.

TTN RNAscope/Iba1 immunofluorescence imaging identified TTN transcripts in both nuclei and cytoplasm of Iba1^+^ cells (Figure 4C). Quantitative analysis indicated that TTN^+^ macrophages were present in control ChP but substantially expanded in AD tissue (Figure 4D), validating the transcriptomic findings of disease-associated expansion of the TTN^+^ macrophage population.

Because many TTN-associated pathways intersect with cellular aging, we also quantified senescence scores across ChP cell types. AD samples exhibited significantly elevated senescence signatures, with macrophages demonstrating the most pronounced increase (Figure 4E). Finally, ligand–receptor inference using LIANA consensus scoring of the ChP snRNA-seq data revealed a broad collapse of macrophage-mediated communication in AD, with reductions in interactions involving epithelial, endothelial, and stromal partners (Figure 4F). Many of the ligand–receptor pairs lost in AD correspond to pathways governing leukocyte recruitment, tissue surveillance, and homeostatic crosstalk, suggesting that disease-associated macrophages become progressively uncoupled from their niche. This broad reduction in intercellular signaling is consistent with the emergence of a TTN^+^, mechanically altered, and senescent macrophage state, which may impair the coordination of immune and barrier functions at the choroid plexus.

## 3 Discussion

The choroid plexus (ChP) is increasingly recognized as a dynamic neuroimmune interface whose cellular states adapt to age, injury, and neurodegeneration. ChP macrophages are a mixed population of long-lived, yolk-sac–derived macrophages and replenishing bone-marrow–derived macrophages that is adapted to the epithelial–stromal interface, positioned at the front line of blood–CSF communication^26,27^. In this study we identified a transcriptionally diverse macrophage compartment within the human ChP. Cross-tissue atlas mapping positions these cells at the intersection of tissue-resident macrophage programs, monocyte-like states, and stress-responsive immune signatures. These findings align with previous descriptions of BAM heterogeneity and niche-driven specialization, while highlighting a broad range of phenotypic states in human ChP macrophages. The background stromal, epithelial, and endothelial cell milieu (Figures 1A–B) underscores the unique structural context in which ChP macrophages operate, supporting immune sampling, antigen presentation, and barrier-associated surveillance.

### ChP macrophages show specialization for their tissue microenvironment

By combining single-nucleus and spatial transcriptomics, our study provides a comprehensive transcriptional and spatial atlas of macrophages in the adult human ChP. BAMs, of which ChP macrophages are a specialized subset, are myeloid cells that reside at central nervous system interfaces, including the meninges and perivascular spaces, where they form a distinct compartment from parenchymal microglia^28^. BAMs are defined by transcriptional programs that integrate immune surveillance, antigen presentation, and regulation of leukocyte trafficking^29,27^. Developmentally, many BAM populations arise from embryonic progenitors and are maintained through local self-renewal^3^, although monocyte-derived cells contribute under inflammatory or disease conditions^26^. Blood-derived macrophages have been demonstrated to play a critical role in CNS tissue repair after injury, mediated in part by IL-10^30,31^. Functionally, BAMs coordinate communication between the peripheral immune system and the CNS, shaping immune entry, inflammatory tone, and tissue homeostasis at neurovascular interfaces, and are increasingly implicated in neurodegenerative and neuroinflammatory diseases.

Among ChP macrophages, three distinct subsets have been described in mice: an Slc40a1^+^ subset located near larger vessels co-expressing Spic and Clec4n; a subset of Lyve1^+^ cells found on the epiplexus surface; and an Spp1 expressing subset^24^. We find subsets overlapping with the three seen in mice. Overlapping with the LYVE1^+^ subset, our data reveals an unexpected mechanobiological program centered on titin (TTN). The discovery of TTN-expressing (TTN^+^) macrophages in the human ChP, and their selective expansion and molecular reprogramming in Alzheimer’s disease (AD), defines a previously unrecognized axis of macrophage mechanotransduction at the brain’s borders.

### Titin expression identifies a distinct mechanosensitive macrophage state

TTN is canonically understood as a structural and elastic scaffold that governs sarcomere mechanics in striated muscle. However, emerging work in lymphocytes demonstrates that immune cells can deploy TTN isoforms to regulate membrane tension, adhesion, and migratory behavior^11,32^. As in other leukocytes^33^, these functions depend on dynamically tuning actin architecture in response to mechanical cues^34^. The identification of TTN transcripts across multiple primer-defined regions in CD163-positive macrophages, together with in situ detection, establishes TTN as a bona fide feature of a macrophage subset in the human ChP. To our knowledge, this represents the first report of TTN expression in a myeloid population in situ, extending TTN biology beyond lymphocyte trafficking and adding to our understanding of immune mechanobiology.

The transcriptional programs enriched in TTN^+^ macrophages including cytoskeletal remodeling, phagocytosis, autophagy–lysosomal activity, stress responses, antigen presentation, and senescence, suggest a functional profile attuned to sustained mechanical load, high turnover demands, and complex interactions within the folded ChP architecture. Interestingly, spatial transcriptomic analysis identified enrichment of the TTN^+^ macrophage population localized near endothelial cells, suggesting a preferential perivascular distribution. Moreover, TTN^+^ cells show heterogeneous but elevated expression of MEF2A, MEF2C, and MEF2D, transcription factors activated downstream of cytoskeletal deformation, Ca^2^+ flux, and RhoA signaling. In proinflammatory conditions, MEF2C contributes to classical activation in macrophages^35^, in contrast to brain microglia, in which it mitigates proinflammatory TNF and IL-1*β* production^36^.

Like other BAMs, ChP macrophages exhibit specialized morphologies and dynamic processes shaped by the mechanical and biochemical attributes of their niche^37^. TTN expression provides a plausible molecular mechanism underlying these specialized cytoskeletal behaviors. The presence of TTN^+^ macrophages at a steady state fills a gap in our understanding of how ChP macrophages sense and respond to mechanical forces within the epithelial–stromal microenvironment. These features position TTN not merely as a marker, but as a potential organizing node, integrating mechanical inputs with transcriptional adaptation in macrophages at the blood–CSF interface.

### Signaling interactions with titin in cytoskeletal control and environmental sensing

Titin has well-characterized interactions with actin and actin-associated complexes—including binding via its PEVK region and Ig-like domains—that enable it to modulate cytoskeletal tension and contribute to force transmission and mechanosensing in cytoskeletal networks under load^38,39^. In lymphocytes, titin is required for mediation of chemokine signalling by RhoA and Rac1^11^. In macrophages, TTN expression may similarly tune the sensitivity of Rho–actin circuits, with downstream effects on force generation, motility, and immune synapse formation. Differential expression and pathway enrichment analyses revealed substantial perturbation of Rho-family small GTPase pathways in AD macrophages. RhoA, Rac1, and Cdc42 orchestrate actin polymerization, contractility, and membrane dynamics that enable leukocyte migration, adhesion, and phagocytosis^40^. Prior studies demonstrate that Rho-GTPase signaling controls macrophage polarity and migration^40,41,42^ and interfaces with inflammatory signaling including NF-*κ*B and oxidative stress responses^43^.

In this framework, TTN^+^ macrophages may be better equipped to navigate the biomechanically complex epithelial folds, fluid shear forces, and variable substrate stiffness of the ChP. The observation that TTN^+^ macrophages occupy spatial niches enriched for endothelial interfaces reinforces the idea that they inhabit a mechanically demanding microenvironment that may selectively engage titin-associated mechanotransduction pathways.

### A disease-associated TTN program emerges in Alzheimer’s disease

Alzheimer’s dis-ease (AD) is increasingly recognized as a disorder involving complex neuroimmune imbalances^13,44,45,46^. While microglial activation in the parenchyma has been extensively studied^15,14^, the choroid plexus (ChP) represents an overlooked interface where peripheral immune cells interact with CNS-derived signals^1,16,17^. The ChP epithelium and its resident macrophages orchestrate leukocyte trafficking into cerebrospinal fluid and regulate inflammatory tone throughout the brain^47,18^. Emerging data indicate that ChP dysfunction may contribute to impaired clearance of toxic proteins and persistent inflammation in AD.

The expansion of TTN^+^ macrophages in AD, supported by both transcriptomic and RNAscope quantification, suggests that this mechanosensitive program becomes preferentially induced or selected under disease conditions. AD macrophages exhibit coordinated upregulation of TTN itself and MEF2-linked signatures, along with Rho-GTPase pathway dysregulation, oxidative stress responses, and hallmarks of cellular senescence. These changes are accompanied by marked downregulation of adhesion- and cytoskeleton-associated transcripts such as RBFOX1, PTPRD, DLG2, and CSMD1, consistent with altered structural and signaling capacity.

A consequence of these transcriptional alterations is the collapse of macrophage interaction networks. Ligand–receptor inference demonstrated that AD macrophages lose connectivity to epithelial, endothelial, and mesenchymal partners. These cells scaffold ChP barrier function and influence leukocyte education. This reduction in communication aligns with the emergence of a TTN^+^, senescent, and spatially segregated macrophage population that may become progressively uncoupled from homeostatic cues. Given the importance of ChP–macrophage interactions for leukocyte recruitment, antigen presentation, and CSF surveillance^48^, such disruption may contribute to impaired waste clearance and amplified inflammatory signaling in AD.

### Implications for neuroimmune biology and future directions

This study provides the first single cell and spatial transcriptomic atlases of adult human choroid plexus from subjects with and without AD, a resource for future studies of the blood-CSF barrier in neurodegeneration. Additionally, our findings introduce titin as a previously unrecognized component of macrophage biology and identify a mechanobiologically specialized macrophage program at the human brain’s borders. By demonstrating that TTN^+^ macrophages display distinct transcriptional signatures, spatial organization, and disease-associated remodeling, this study expands the conceptual framework of immune mechanotransduction and implicates TTN in neuroimmune dysfunction.

Future work should determine whether TTN directly regulates macrophage mechanics, migration, or adhesion in the ChP; whether specific TTN isoforms confer specialized biomechanical properties; and whether TTN-associated pathways can be therapeutically modulated to restore macrophage–epithelial crosstalk in AD. More broadly, these results raise the possibility that mechanical defects at CNS interfaces, rather than solely in parenchymal tissue, contribute to the immune dysregulation characteristic of Alzheimer’s disease.

## 4 Methods

### 4.1 Choroid Plexus Tissue Preparation

Choroid plexus tissues were dissected from the bilateral lateral ventricles of deidentifed post mortem human donors with PMI *<*6 hours through the Yale Legacy Tissue Donation Program. ChP samples were divided, with portions of tissue *>*10 grams snap frozen and a portion fixed in 10% formalin and embedded in paraffin. Donor metadata is provided in Supplementary Materials Table S1.

### 4.2 Single-Nucleus RNA Sequencing

Nuclei were isolated on ice using the Nuclei EZ Prep Kit (NUC-101, Sigma-Aldrich) according to the manufacturer’s instructions, with some modification. All procedures were carried out on ice or at 4°C. Briefly, frozen tissue was subjected to dounce homogenization (25 times with pestle A followed by 25 times with tight pestle B) using the KIMBLE Dounce Tissue Grinder Set (Sigma) in 2ml EZ Lysis buffer. The sample was transferred to a 15ml tube with an additional 2ml EZ lysis buffer and incubated on ice for 5 minutes. Following incubation, the sample was centrifuged at 500xg for 5 minutes at 4°C. Supernatants were discarded, and the isolated nuclei were resuspended in 4ml EZ lysis buffer, incubated for 5 minutes on ice and centrifuged at 500xg for 5 minutes at 4°C. Next, the nuclei were washed with 4ml ice-cold Nuclei Suspension Buffer (1X PBS containing 0.01% BSA and 0.1% RNAse inhibitor), resuspended in 1ml Nuclei EZ Storage buffer and passed through a 40 µm nylon cell strainer. The nuclei suspensions were counted with trypan blue using a hemocytometer under a microscope. RNA libraries of barcoded single nuclei were assembled in compliance with the 10X Chromium Next GEM Single Cell 3’ platforms manufacturer procedures. Sequencing was performed by the Yale Center for Genome Analysis (YCGA) using an Illumina NovaSeq 6000, ensuring high-quality single nucleus RNA sequencing data for downstream analysis. The raw data obtained from sequencing was converted to fastq files, which were then aligned to the human reference genome (GRCh38) using the 10X Cell Ranger 7.1.0 pipeline.

### 4.3 Spatial transcriptomics

OCT-embedded snap frozen tissues were cryosectioned at 10 µm thickness and applied to barcoded Curio Bioscience Seeker v1.1 3×3mm slide-seq tiles according to the manufacturer’s protocol. Sections were fixed, permeabilized, and processed for spatially resolved cDNA synthesis using Slide-seq reagents. Libraries were prepared and sequenced using an Illumina NovaSeq 6000, achieving a mean spatial resolution of 10 µm per bead. Raw sequencing data were processed using the Curio computational pipeline implemented in Nextflow. Spatial barcode deconvolution, gene count matrices, and spatial alignment were performed using the Curio pipeline and validated with gene signatures and anatomical landmarks. After batch correction with Harmony^49^, we used GIMVI^19^ to integrate into the spatial transcriptome dataset expression data our snRNA-seq dataset from ChP to increase gene coverage. We then used CellCharter^50^ to perform graph-based coarse neighborhood clustering, and identified cell types using gene signature scores for canonical cell-type marker sets, assigning each cell to the class whose marker signature achieved the maximal score for that cell.

### 4.4 CD163^+^ Cell Selection, RNA Extraction, and cDNA Synthesis

RNA was extracted from CD163 microbead-separated macrophages (Miltenyi Biotec, Cat. #130-124-420) using the RNeasy Mini Kit (Qiagen), following the manufacturer’s protocol. The extracted RNA was then quantified, and its purity and integrity were assessed using a NanoDrop 2000 spectrophotometer (Thermo Scientific). For cDNA synthesis, 12 ng of RNA-equivalent cDNA was generated. The cDNA synthesis was performed using the iScript cDNA Synthesis Kit (Bio-Rad, Cat. 1708891), ensuring high-quality template for subsequent qPCR analysis. The cDNA was subsequently amplified using TTN-specific primers (IDT DNA; For sequences of primers used, see Supplementary Table S5).

### 4.5 Validation and Analysis of cDNA Using qPCR

The TTN expression levels were assessed using quantitative PCR (qPCR) conducted with a Bio-Rad system. The generated cDNA was validated using qPCR, and the amplification plots were analyzed with Bio-Rad CFX Maestro software. This analysis provided detailed amplification profiles of the expression of TTN in the CD163 microbead-separated macrophages, with GAPDH serving as a control to ensure the specificity and reliability of the assay.

### 4.6 Immunofluorescence and RNA in situ hybridization

Formalin-fixed, paraffin-embedded (FFPE) tissue blocks were sectioned into 5µm thick slices and mounted on Superfrost Plus microscope slides. The slides were baked overnight at 40°C to ensure adherence of the sections, deparaffinized, and hydrated prior to staining.

For titin immunofluorescence, primary antibody incubation with rabbit anti-titin (Proteintech Catalog #27867-1-AP) at 1:250 dilution and mouse anti-CD68 antibody (Millipore Cat.#168M-95) at 1:250 was conducted overnight at 4°C, followed by detection with secondary donkey anti-rabbit Alexa Fluor 488 (Thermo Fisher Scientific, Cat. #A21206) (1:500) and goat anti-mouse Alexa Fluor 647 (Thermo Fisher Scientific, Cat. #A32728), with nuclei counterstaining with DAPI.

RNA probes were obtained from ACDBio, Inc., Newark, CA, for TTN (RNAscope Probe Hs-TTN, Cat. #550361) and detected using TSA Vivid Fluorophore Kit 650 (Tocris Bioscience, Cat. #7527). RNAscope was performed following manufacturer’s protocol. Primary antibody incubation with Rabbit Anti-Iba1 (Abcam Inc., Cat. #ab178847) at 1:250 dilution in 1% BSA with 0.1% Tween-20 was performed overnight at 4°C. Secondary detection used Donkey anti-Rabbit IgG (H+L) Highly Cross-Adsorbed Secondary Antibody, Alexa Fluor 488 (Thermo Fisher Scientific, Cat. #A21206) (1:500), with nuclei counterstaining with DAPI.

Slides were imaged on an inverted fluorescence microscope (Thunder Imager, Leica Microsystems, Wetzlar, Germany). A minimum of three slides were imaged per condition, with a mean of 27.2 images per slide and a mean of 24.4 images per case. The total number of macrophages and the number of macrophages with probe signal in areas of 558.14 µm x 348.84 µm were recorded and pooled based on condition (AD or control). A total of 310 images were analyzed: 91 images from control slides (n=2) and 221 images from AD slides (n=7). Statistical analyses were performed using a Bayesian independent sample t-test (H*_a_*= AD*>*Control).

### 4.7 DAB Immunohistochemistry

FFPE sections were deparaffinized in xylene and rehydrated using graded alcohols. Antigen retrieval was performed using citrate buffer, followed by blocking with 1% BSA and 0.3% Triton X-100. Primary antibody incubation with rabbit anti-CD163 (Abcam Inc., Cat. #ab182422) at 1:200 dilution was conducted overnight at 4°C. Detection involved biotinylated goat anti-rabbit IgG and ABC reagent, with visualization via DAB substrate.

### 4.8 Computational Analysis

The snRNA-seq data were analyzed using Python-based tools, including scVI, Scanpy, and GSEA, for statistical analysis and visualization. Additional custom Python code was used for spatial ROI selection and analysis (see Data Availability section).

#### Cell Composition Analysis

To characterize the baseline cellular architecture of the tissue independent of experimental perturbation, we examined the intercept terms of a scCODA model fitted to our data, which represent the log-scale baseline abundances of each cell type relative to the reference population. scCODA implements a Bayesian multinomial–logit framework with a spike-and-slab prior, allowing simultaneous estimation of positive, negative, or null compositional effects while accounting for the inherent sum-to-one constraints of compositional data. By transforming the posterior mean intercepts through a softmax function, we derived an interpretable baseline composition profile reflecting the expected proportion of each cell type before conditioning on the experimental variable. This intercept-derived composition provides a model-based estimate of the underlying cellular milieu, integrating information across all samples while adjusting for reference constraints and compositional dependencies. In our dataset, the resulting stacked bar representation highlights the baseline dominance of specific populations and provides a principled point of comparison for evaluating subsequent condition-associated shifts in relative abundance.

Gene module scores were computed using scanpy.tl.score genes, which measures the average expression of a predefined gene set in each cell relative to a background reference set of matched genes. Scores are centered around zero, such that positive values indicate enrichment of the module and negative values indicate depletion relative to background. Gene sets for scoring were curated from a combination of publicly available pathway databases (MSigDB, Reactome), ImmGen and human myeloid single-cell atlases, and manual literature curation. Specific genes used for myeloid cell types (Figure 1D) are listed in Supplementary Table S3, and genes used for functional annotations (Figure 3A) are listed in Supplementary Table S4.

#### Atlas integration

To contextualize choroid plexus macrophages within a broader myeloid landscape, we performed cross-dataset integration between our macrophage snRNA-seq data and the myeloid compartment of a cross tissue immune cell atlas^23^ using a non-negative matrix factorization (NMF) framework implemented in python with the scikit-learn library. NMF decomposes high-dimensional single-nucleus gene expression matrices into a set of biologically interpretable latent factors whose non-negativity constraints promote sparse, additive representations of cellular programs. For integration, we projected both our choroid plexus dataset and external reference atlases of human myeloid populations into a shared NMF-derived factor space. This approach aligns gene expression profiles across datasets without imposing hard cluster boundaries and enables robust comparison of transcriptional identities across tissues. The resulting joint factorization allowed us to map choroid plexus macrophages onto well-defined myeloid states.

#### Differential Gene Expression

To define transcriptional changes between conditions and macrophage subpopulations, we performed differential gene expression (DGE) analysis on scVI-corrected expression values. Statistical testing relied on a generalized linear modeling approach with empirical Bayes variance moderation to account for heteroscedasticity inherent to single-nucleus data. Genes were considered significantly differentially expressed if they satisfied an adjusted p-value less than 0.05 and an absolute fold change greater than 50.

#### Spatial Neighborhood Analysis

To integrate transcriptional states with spatial organization in the choroid plexus, we performed a multimodal spatial neighborhood analysis combining cross-modal imputation, graph-based clustering, and neighborhood enrichment statistics. First, we used GIMVI^19^, a variational inference framework that jointly models snRNA-seq and spatial transcriptomics data, to impute high-resolution gene expression profiles into Visium grid locations. This approach enables transfer of macrophage subcluster signatures—such as TTN-positive programs—onto the tissue architecture while preserving spatial covariance across modalities. We then applied CellCharter^50^, which performs graph-based clustering on spatial expression graphs, to delineate coherent anatomical and functional domains within the choroid plexus. These domains captured epithelial folds, stromal regions, and immune-enriched microenvironments and provided a spatial scaffold for downstream analyses. Finally, using Squidpy’s neighborhood enrichment framework^51^, we quantified the overrepresentation of specific cell types in the immediate proximity of TTN-positive and TTN-negative macrophages.

#### Gene Set Enrichment Analysis and Senescence scoring

To investigate coordinated pathway-level changes, we performed gene set enrichment analysis (GSEA) using Enrichr^52^ across Gene Ontology Biological Process, KEGG, and Reactome databases. Enrichment statistics were derived from the ranked DGE results. To quantitatively assess cellular aging programs, we used senepy^53^, a framework for single-cell resolution senescence scoring. Senepy computes enrichment scores for curated senescence gene sets while adjusting for dataset-specific expression variability.

#### Cell Interaction Analysis

Cell-cell communication was inferred using LIANA^54^, which provides consensus scoring across multiple ligand-receptor inference algorithms to ensure robustness to model-specific biases. We applied LIANA to scVI-normalized expression profiles in our snRNAseq data to quantify the probability of active ligand-receptor interactions between all annotated cell classes. Consensus scores were aggregated and visualized using circos plots to highlight directional communication patterns in control and Alzheimer’s disease tissue.

## 5 Data availability

The snRNA-seq and spatial transcriptomics data generated in this study have been deposited in the NCBI Gene Expression Omnibus (GEO) under accession numbers GSE315551 and GSE315553, respectively. Processed count matrices, cell annotations, and metadata used for all analyses are included in the GEO record. Additional data supporting the findings of this study are available from the corresponding author upon reasonable request. Custom analysis code is available at https://github.com/distasiolab/SHiPBIO.

## Acknowledgments

We would like to thank the tissue donors and their families for their contribution to this work through the Yale Legacy Tissue Donation Program. Without their generosity, our study would not have been possible. M.D. receives research funding from NEI K08-EY033013.

## 6 Author Contributions

Conception: M.D.; Design of Work: M.D., S.B., A.G.; Acquisition of Data: M.D., S.B., A.G., G.O.; Analysis of Data: M.D., S.B., A.G., G.O.; Interpretation of Data: M.D.; Writing and Editing: M.D., S.B., A.G., G.O.

## References

[1] H. Xu, P. Lotfy, S. Gelb, et al. “The choroid plexus synergizes with immune cells during neuroinflammation”. English. In: Cell 187.18 (Sept. 2024). Publisher: Elsevier, 4946–4963.e17. ISSN: 0092-8674, 1097-4172. DOI: 10.1016/j.cell.2024.07.002. URL: https://www.cell.com/cell/abstract/S0092-8674(24)00717-7 (visited on 12/03/2024).

[2] H. Van Hove, C. Glü ck, W. Mildenberger, et al. “Interleukin-34-dependent perivascular macrophages promote vascular function in the brain”. In: Immunity 58.5 (May 2025), 1289–1305.e8. ISSN: 1074-7613. DOI: 10.1016/j.immuni.2025.04.003. URL: https://www.sciencedirect.com/science/article/pii/S1074761325001669 (visited on 09/18/2025).

[3] S. G. Utz, P. See, W. Mildenberger, et al. “Early Fate Defines Microglia and Non-parenchymal Brain Macrophage Development”. English. In: Cell 181.3 (Apr. 2020). Publisher: Elsevier, 557–573.e18. ISSN: 0092-8674, 1097-4172. DOI: 10.1016/j.cell.2020.03.021. URL: https://www.cell.com/cell/abstract/S0092-8674(20)30283-X (visited on 11/18/2025).

[4] M. Guilliams, G. R. Thierry, J. Bonnardel, et al. “Establishment and Maintenance of the Macrophage Niche”. en. In: Immunity 52.3 (Mar. 2020), pp. 434–451. ISSN: 10747613. DOI: 10.1016/j.immuni.2020.02.015. URL: https://linkinghub.elsevier.com/retrieve/pii/S1074761320300868 (visited on 09/18/2025).

[5] H. Du, J. M. Bartleson, S. Butenko, et al. “Tuning immunity through tissue mechanotransduction”. en. In: Nature Reviews Immunology 23.3 (Mar. 2023). Publisher: Nature Publishing Group, pp. 174–188. ISSN: 1474-1741. DOI: 10.1038/s41577-022-00761-w. URL: https://www.nature.com/articles/s41577-022-00761-w (visited on 12/13/2025).

[6] P. W. Oakes, E. Wagner, C. A. Brand, et al. “Optogenetic control of RhoA reveals zyxin-mediated elasticity of stress fibres”. eng. In: Nature Communications 8 (June 2017), p. 15817. ISSN: 2041-1723. DOI: 10.1038/ncomms15817.

[7] M. L. Dustin and J. A. Cooper. “The immunological synapse and the actin cytoskeleton: molecular hardware for T cell signaling”. en. In: Nature Immunology 1.1 (July 2000). Publisher: Nature Publishing Group, pp. 23–29. ISSN: 1529-2916. DOI: 10.1038/76877. URL: https://www.nature.com/articles/ni0700_23 (visited on 12/13/2025).

[8] F. Y. McWhorter, T. Wang, P. Nguyen, et al. “Modulation of macrophage phenotype by cell shape”. In: Proceedings of the National Academy of Sciences 110.43 (Oct. 2013). Publisher: Proceedings of the National Academy of Sciences, pp. 17253–17258. DOI: 10.1073/pnas.1308887110. URL: https://www.pnas.org/doi/10.1073/pnas.1308887110 (visited on 12/13/2025).

[9] X. Zhang, T.-H. Kim, T. J. Thauland, et al. “Unraveling the mechanobiology of immune cells”. In: Current Opinion in Biotechnology 66 (Dec. 2020), pp. 236–245. ISSN: 0958-1669. DOI: 10.1016/j.copbio.2020.09.004. URL: https://pmc.ncbi.nlm.nih.gov/articles/PMC7524653/ (visited on 12/16/2025).

[10] S. Labeit, B. Kolmerer, and W. A. Linke. “The giant protein titin. Emerging roles in physiology and pathophysiology”. eng. In: Circulation Research 80.2 (Feb. 1997), pp. 290–294. ISSN: 0009-7330. DOI: 10.1161/01.res.80.2.290.

[11] L. Toffali, B. D’Ulivo, C. Giagulli, et al. “An isoform of the giant protein titin is a master regulator of human T lymphocyte trafficking”. eng. In: Cell Reports 42.5 (May 2023), p. 112516. ISSN: 2211-1247. DOI: 10.1016/j.celrep.2023.112516.

[12] A. Mammoto, T. Mammoto, and D. E. Ingber. “Mechanosensitive mechanisms in transcriptional regulation”. In: Journal of Cell Science 125.13 (July 2012), pp. 3061–3073. ISSN: 0021-9533. DOI: 10.1242/jcs.093005. URL: https://pmc.ncbi.nlm.nih.gov/articles/PMC3434847/ (visited on 12/13/2025).

[13] M. T. Heneka, W. M. van der Flier, F. Jessen, et al. “Neuroinflammation in Alzheimer disease”. en. In: Nature Reviews Immunology 25.5 (May 2025). Publisher: Nature Publishing Group, pp. 321–352. ISSN: 1474-1741. DOI: 10.1038/s41577-024-01104-7. URL: https://www.nature.com/articles/s41577-024-01104-7 (visited on 12/12/2025).

[14] D. V. Hansen, J. E. Hanson, and M. Sheng. “Microglia in Alzheimer’s disease”. In: The Journal of Cell Biology 217.2 (Feb. 2018), pp. 459–472. ISSN: 0021-9525. DOI: 10.1083/jcb.201709069. URL: https://pmc.ncbi.nlm.nih.gov/articles/PMC5800817/ (visited on 12/12/2025).

[15] S. Hong, V. F. Beja-Glasser, B. M. Nfonoyim, et al. “Complement and microglia mediate early synapse loss in Alzheimer mouse models”. In: Science 352.6286 (May 2016). Publisher: American Association for the Advancement of Science, pp. 712–716. DOI: 10.1126/science.aad8373. URL: https://www.science.org/doi/10.1126/science.aad8373 (visited on 12/12/2025).

16. I. Strominger, Y. Elyahu, O. Berner, et al. “The Choroid Plexus Functions as a Niche for T-Cell Stimulation Within the Central Nervous System”. In: Frontiers in Immunology 9 (2018). ISSN: 1664-3224. URL: https://www.frontiersin.org/journals/immunology/articles/10.3389/fimmu.2018.01066 (visited on 02/22/2024).

[17] K. Baruch, A. Deczkowska, E. David, et al. “Aging-induced type I interferon response at the choroid plexus negatively affects brain function”. In: Science 346.6205 (Oct. 2014). Publisher: American Association for the Advancement of Science, pp. 89–93. DOI: 10.1126/science.1252945. URL: https://www.science.org/doi/10.1126/science.1252945 (visited on 01/08/2025).

[18] R. Shechter, A. London, and M. Schwartz. “Orchestrated leukocyte recruitment to immune-privileged sites: absolute barriers versus educational gates”. en. In: Nature Reviews Immunology 13.3 (Mar. 2013). Publisher: Nature Publishing Group, pp. 206–218. ISSN: 1474-1741. DOI: 10.1038/nri3391. URL: https://www.nature.com/articles/nri3391 (visited on 12/12/2025).

[19] R. Lopez, A. Nazaret, M. Langevin, et al. “A joint model of unpaired data from scRNA-seq and spatial transcriptomics for imputing missing gene expression measurements”. In: *ArXiv* (May 2019). URL: https://www.semanticscholar.org/paper/A-joint-model-of-unpaired-data-from-scRNA-seq-and-Lopez-Nazaret/9991c277fd6a0e2363631846e9a7ea40abc9b7dc (visited on 12/15/2025).

[20] F. A. Wolf, P. Angerer, and F. J. Theis. “SCANPY: large-scale single-cell gene expression data analysis”. eng. In: Genome Biology 19.1 (Feb. 2018), p. 15. ISSN: 1474-760X. DOI: 10.1186/s13059-017-1382-0.

[21] M. Bü ttner, J. Ostner, C. L. Mü ller, et al. “scCODA is a Bayesian model for compositional single-cell data analysis”. en. In: Nature Communications 12.1 (Nov. 2021). Publisher: Nature Publishing Group, p. 6876. ISSN: 2041-1723. DOI: 10.1038/s41467-021-27150-6. URL: https://www.nature.com/articles/s41467-021-27150-6 (visited on 12/13/2025).

[22] I. Tirosh, B. Izar, S. M. Prakadan, et al. “Dissecting the multicellular ecosystem of metastatic melanoma by single-cell RNA-seq”. eng. In: Science (New York, N.Y.) 352.6282 (Apr. 2016), pp. 189–196. ISSN: 1095-9203. DOI: 10.1126/science.aad0501.

[23] C. Domínguez Conde, C. Xu, L. B. Jarvis, et al. “Cross-tissue immune cell analysis reveals tissue-specific features in humans”. In: Science 376.6594 (May 2022). Publisher: American Association for the Advancement of Science, eabl5197. DOI: 10.1126/science.abl5197. URL: https://www.science.org/doi/10.1126/science.abl5197 (visited on 11/13/2025).

[24] N. Dani, R. H. Herbst, C. McCabe, et al. “A cellular and spatial map of the choroid plexus across brain ventricles and ages”. eng. In: Cell 184.11 (May 2021), 3056–3074.e21. ISSN: 1097-4172. DOI: 10.1016/j.cell.2021.04.003.

[25] S. G. Rodriques, R. R. Stickels, A. Goeva, et al. “Slide-seq: A scalable technology for measuring genome-wide expression at high spatial resolution”. In: Science 363.6434 (Mar. 2019). Publisher: American Association for the Advancement of Science, pp. 1463–1467. DOI: 10.1126/science.aaw1219. URL: https://www.science.org/doi/10.1126/science.aaw1219 (visited on 12/18/2025).

[26] T. Goldmann, P. Wieghofer, M. J. C. Jordão, et al. “Origin, fate and dynamics of macrophages at central nervous system interfaces”. en. In: Nature Immunology 17.7 (July 2016). Publisher: Nature Publishing Group, pp. 797–805. ISSN: 1529-2916. DOI: 10.1038/ni.3423. URL: https://www.nature.com/articles/ni.3423 (visited on 01/02/2026).

[27] H. Van Hove, L. Martens, I. Scheyltjens, et al. “A single-cell atlas of mouse brain macrophages reveals unique transcriptional identities shaped by ontogeny and tissue environment”. eng. In: Nature Neuroscience 22.6 (June 2019), pp. 1021–1035. ISSN: 1546-1726. DOI: 10.1038/s41593-019-0393-4.

[28] A. Zeisel, A. B. Muñoz-Manchado, S. Codeluppi, et al. “Cell types in the mouse cortex and hippocampus revealed by single-cell RNA-seq”. In: Science 347.6226 (Mar. 2015). Publisher: American Association for the Advancement of Science, pp. 1138–1142. DOI: 10.1126/science.aaa1934. URL: https://www.science.org/doi/10.1126/science.aaa1934 (visited on 12/17/2025).

[29] D. Mrdjen, A. Pavlovic, F. J. Hartmann, et al. “High-Dimensional Single-Cell Mapping of Central Nervous System Immune Cells Reveals Distinct Myeloid Subsets in Health, Aging, and Disease”. eng. In: Immunity 48.2 (Feb. 2018), 380–395.e6. ISSN: 1097-4180. DOI: 10.1016/j.immuni.2018.01.011.

[30] R. Shechter, A. London, C. Varol, et al. “Infiltrating Blood-Derived Macrophages Are Vital Cells Playing an Anti-inflammatory Role in Recovery from Spinal Cord Injury in Mice”. en. In: PLOS Medicine 6.7 (July 2009). Publisher: Public Library of Science, e1000113. ISSN: 1549-1676. DOI: 10.1371/journal.pmed.1000113. URL: https://journals.plos.org/plosmedicine/article?id=10.1371/journal.pmed.1000113 (visited on 01/05/2026).

[31] A. London, E. Itskovich, I. Benhar, et al. “Neuroprotection and progenitor cell renewal in the injured adult murine retina requires healing monocyte-derived macrophages”. In: Journal of Experimental Medicine 208.1 (Jan. 2011), pp. 23–39. ISSN: 0022-1007. DOI: 10.1084/jem.20101202. (visited on 01/05/2026).

[32] B. J. de Andrade Silva, T. K. Hughes, P. R. Andrade, et al. “Single-cell RNA-sequencing of leprosy skin lesions reveals a titin (TTN) expressing T cell subset associated with the progressive disease”. In: The Journal of Immunology 204.1 Supplement (May 2020), p. 225.36. ISSN: 0022-1767. URL: 10.4049/jimmunol.204.Supp.225.36 (visited on 11/05/2025).

[33] T. Lä mmermann and R. N. Germain. “The multiple faces of leukocyte interstitial migration”. eng. In: Seminars in Immunopathology 36.2 (Mar. 2014), pp. 227–251. ISSN: 1863-2300. DOI: 10.1007/s00281-014-0418-8.

[34] F. Y. McWhorter, C. T. Davis, and W. F. Liu. “Physical and mechanical regulation of macrophage phenotype and function”. In: Cellular and Molecular Life Sciences: CMLS 72.7 (Dec. 2014), pp. 1303–1316. ISSN: 1420-682X. DOI: 10.1007/s00018-014-1796-8. URL: https://pmc.ncbi.nlm.nih.gov/articles/PMC4795453/ (visited on 01/05/2026).

[35] X. Zhao, Q. Di, H. Liu, et al. “MEF2C promotes M1 macrophage polarization and Th1 responses”. en. In: Cellular & Molecular Immunology 19.4 (Apr. 2022). Publisher: Nature Publishing Group, pp. 540–553. ISSN: 2042-0226. DOI: 10.1038/s41423-022-00841-w. URL: https://www.nature.com/articles/s41423-022-00841-w (visited on 01/05/2026).

[36] A. Deczkowska, O. Matcovitch-Natan, A. Tsitsou-Kampeli, et al. “Mef2C restrains microglial inflammatory response and is lost in brain ageing in an IFN-I-dependent manner”. en. In: Nature Communications 8.1 (Sept. 2017). Publisher: Nature Publishing Group, p. 717. ISSN: 2041-1723. DOI: 10.1038/s41467-017-00769-0. URL: https://www.nature.com/articles/s41467-017-00769-0 (visited on 01/05/2026).

[37] K. Kierdorf, T. Masuda, M. J. C. Jordão, et al. “Macrophages at CNS interfaces: ontogeny and function in health and disease”. en. In: Nature Reviews Neuroscience 20.9 (Sept. 2019). Publisher: Nature Publishing Group, pp. 547–562. ISSN: 1471-0048. DOI: 10. 1038/s41583-019-0201-x. URL: https://www.nature.com/articles/s41583-019-0201-x (visited on 12/16/2025).

[38] P. Bianco, A. Nagy, A. Kengyel, et al. “Interaction Forces between F-Actin and Titin PEVK Domain Measured with Optical Tweezers”. English. In: Biophysical Journal 93.6 (Sept. 2007). Publisher: Elsevier, pp. 2102–2109. ISSN: 0006-3495, 1542-0086. DOI: 10.1529/biophysj.107.106153. URL: https://www.cell.com/biophysj/abstract/S0006-3495(07)71464-9 (visited on 12/18/2025).

[39] C. M. Loescher, A. J. Hobbach, and W. A. Linke. “Titin (TTN): from molecule to modifications, mechanics, and medical significance”. In: Cardiovascular Research 118.14 (Oct. 2021), pp. 2903–2918. ISSN: 0008-6363. DOI: 10.1093/cvr/cvab328. URL: https://pmc.ncbi.nlm.nih.gov/articles/PMC9648829/ (visited on 12/18/2025).

[40] W. E. Allen, G. E. Jones, J. W. Pollard, et al. “Rho, Rac and Cdc42 regulate actin organization and cell adhesion in macrophages”. eng. In: Journal of Cell Science 110 ( Pt 6) (Mar. 1997), pp. 707–720. ISSN: 0021-9533. DOI: 10.1242/jcs.110.6.707.

[41] V. Kö nigs, R. Jennings, T. Vogl, et al. “Mouse macrophages completely lacking Rho subfamily GTPases (RhoA, RhoB, and RhoC) have severe lamellipodial retraction defects, but robust chemotactic navigation and altered motility”. eng. In: The Journal of Biological Chemistry 289.44 (Oct. 2014), pp. 30772–30784. ISSN: 1083-351X. DOI: 10.1074/jbc.M114.563270.

[42] V. S. Meli, P. K. Veerasubramanian, H. Atcha, et al. “Biophysical regulation of macrophages in health and disease”. In: Journal of leukocyte biology 106.2 (Aug. 2019), pp. 283–299. ISSN: 0741-5400. DOI: 10.1002/JLB.MR0318-126R. URL: https://pmc.ncbi.nlm.nih.gov/articles/PMC7001617/ (visited on 12/16/2025).

[43] B.-C. Chen, J.-C. Kang, Y.-T. Lu, et al. “Rac1 regulates peptidoglycan-induced nuclear factor-B activation and cyclooxygenase-2 expression in RAW 264.7 macrophages by activating the phosphatidylinositol 3-kinase/Akt pathway”. In: Molecular Immunology 46.6 (Mar. 2009), pp. 1179–1188. ISSN: 0161-5890. DOI: 10.1016/j.molimm.2008.11.006. URL: https://www.sciencedirect.com/science/article/pii/S0161589008007542 (visited on 12/16/2025).

[44] M. Ito, K. Komai, S. Mise-Omata, et al. “Brain regulatory T cells suppress astrogliosis and potentiate neurological recovery”. en. In: Nature 565.7738 (Jan. 2019). Publisher: Nature Publishing Group, pp. 246–250. ISSN: 1476-4687. DOI: 10.1038/s41586-018-0824-5. URL: https://www.nature.com/articles/s41586-018-0824-5 (visited on 12/12/2025).

[45] Y. Zhang, I. T. H. Fung, P. Sankar, et al. “Depletion of nk cells improves cognitive function in the alzheimer disease mouse model”. English. In: Journal of Immunology 205.2 (2020). Publisher: American Association of Immunologists, pp. 502–510. ISSN: 0022-1767. DOI: 10.4049/jimmunol.2000037.

[46] E. Zenaro, E. Pietronigro, V. D. Bianca, et al. “Neutrophils promote Alzheimer’s dis-ease–like pathology and cognitive decline via LFA-1 integrin”. en. In: Nature Medicine 21.8 (Aug. 2015). Publisher: Nature Publishing Group, pp. 880–886. ISSN: 1546-170X. DOI: 10.1038/nm.3913. URL: https://www.nature.com/articles/nm.3913 (visited on 12/12/2025).

[47] R. Shechter, O. Miller, G. Yovel, et al. “Recruitment of beneficial M2 macrophages to injured spinal cord is orchestrated by remote brain choroid plexus”. eng. In: Immunity 38.3 (Mar. 2013), pp. 555–569. ISSN: 1097-4180. DOI: 10.1016/j.immuni.2013.02.012.

[48] M. Schwartz and K. Baruch. “The resolution of neuroinflammation in neurodegeneration: leukocyte recruitment via the choroid plexus”. eng. In: The EMBO journal 33.1 (Jan. 2014), pp. 7–22. ISSN: 1460-2075. DOI: 10.1002/embj.201386609.

[49] I. Korsunsky, N. Millard, J. Fan, et al. “Fast, sensitive and accurate integration of single-cell data with Harmony”. en. In: Nature Methods 16.12 (Dec. 2019). Publisher: Nature Publishing Group, pp. 1289–1296. ISSN: 1548-7105. DOI: 10.1038/s41592-019-0619-0. URL: https://www.nature.com/articles/s41592-019-0619-0 (visited on 12/15/2025).

[50] M. Varrone, D. Tavernari, A. Santamaria-Martínez, et al. “CellCharter reveals spatial cell niches associated with tissue remodeling and cell plasticity”. eng. In: Nature Genetics 56.1 (Jan. 2024), pp. 74–84. ISSN: 1546-1718. DOI: 10.1038/s41588-023-01588-4.

[51] G. Palla, H. Spitzer, M. Klein, et al. “Squidpy: a scalable framework for spatial omics analysis”. en. In: Nature Methods 19.2 (Feb. 2022). Publisher: Nature Publishing Group, pp. 171–178. ISSN: 1548-7105. DOI: 10.1038/s41592-021-01358-2. URL: https://www.nature.com/articles/s41592-021-01358-2 (visited on 12/16/2025).

[52] E. Y. Chen, C. M. Tan, Y. Kou, et al. “Enrichr: interactive and collaborative HTML5 gene list enrichment analysis tool”. eng. In: BMC bioinformatics 14 (Apr. 2013), p. 128. ISSN: 1471-2105. DOI: 10.1186/1471-2105-14-128.

[53] M. A. Sanborn, X. Wang, S. Gao, et al. “Unveiling the cell-type-specific landscape of cellular senescence through single-cell transcriptomics using SenePy”. en. In: Nature Communications 16.1 (Feb. 2025). Publisher: Nature Publishing Group, p. 1884. ISSN: 2041-1723. DOI: 10.1038/s41467-025-57047-7. URL: https://www.nature.com/articles/s41467-025-57047-7 (visited on 12/16/2025).

[54] D. Dimitrov, P. S. L. Schäfer, E. Farr, et al. “LIANA+ provides an all-in-one framework for cell–cell communication inference”. en. In: Nature Cell Biology 26.9 (Sept. 2024). Publisher: Nature Publishing Group, pp. 1613–1622. ISSN: 1476-4679. DOI: 10.1038/s41556-024-01469-w. URL: https://www.nature.com/articles/s41556-024-01469-w (visited on 11/30/2025).

