## Supplemental Information for "Mechanobiological Specialization of Choroid Plexus Macrophages Defined by Titin Expression"

#### SUPPLEMENTARY MATERIALS

Sagar Bhatta[1], Alayna Grzybowski[2], Gion Ortiz[3], and Marcello DiStasio[1,2,*]

[1]Department of Ophthalmology and Visual Science, Yale School of Medicine, New Haven, CT

[2]Department of Pathology Yale School of Medicine, New Haven, CT

[3]Program in Forensic Science, University of New Haven, New Haven, CT

### 1 Supplementary Tables

| Subject | Age | Sex | PMI | Dementia | Pathologic/Clinical History | snRNA seq | RNA scope | Spatial RNAseq | qPCR |
|---|---|---|---|---|---|---|---|---|---|
| A.1 | 36 | M | 11h | (-) | Cerebrovascular disease | ✓ | | | |
| A.2 | 66 | F | 25h | (-) | Squamous cell carcinoma of lung | ✓ | | | |
| A.3 | 78 | M | 23h | (-) | Pneumonia | ✓ | | | |
| A.4 | 68 | F | 32h | (-) | Myocardial infarction | ✓ | | | |
| A.5 | 82 | M | 22h | AD | High level of ADNC (A3,B3,C2) | ✓ | | ✓ | |
| A.6 | 55 | F | 14h | AD | High level of ADNC (A3,B3,C3) | ✓ | | | |
| YA.1 | 69 | F | 12h | (-) | No history of cognitive disorder | ✓ | | | |
| YA.2 | 88 | F | 9h | AD | High level of ADNC (A3,B3,C3) | ✓ | | | |
| YA.3 | 79 | M | 5h | AD | Alzheimer's disease; pathologic stage unknown | ✓ | | | |
| YA.4 | 89 | M | 6h | AD | Intermediate level of ADNC (A1,B3,C2); Pneumonia | | | ✓ | |
| YA.5 | 87 | F | 4h | AD | Intermediate level of ADNC (A3,B2,C2) | | | ✓ | |
| YA.6 | 63 | M | 5h | AD | High level of ADNC (A3,B3,C2); LBD, transitional/limbic | | | ✓ | |
| A.7 | 77 | M | 114h | (-) | Pneumonia | | ✓ | | |
| A.8 | 70 | M | | (-) | Pulmonary embolism | | ✓ | | |
| A.9 | 59 | F | 22h | AD | High level of ADNC (A3,B3,C3) | | ✓ | | |
| A.10 | 88 | F | 9h | AD | High level of ADNC (A3,B3,C3) | | ✓ | | |
| A.11 | 86 | F | 4h | AD | Alzheimer's disease; pathologic stage unknown | | ✓ | | |
| A.12 | 89 | F | 15h | AD | Intermediate level of ADNC (A2,B3,C2); Remote infarct, R. occipital cortex | | ✓ | | |
| A.13 | 84 | F | 41h | AD | Intermediate level of ADNC (A2,B3,C2) | | ✓ | | |
| A.14 | 94 | F | 45h | AD | Low level of ADNC (A1,B2,C2) | | ✓ | | |
| YA.7 | 62 | F | 4h | (-) | No history of cognitive disorder | | | | ChP |
| YA.8 | 84 | F | 18h | AD | Alzheimer's disease; pathologic stage unknown | | | | ChP |
| YA.9 | 88 | M | 6h | AD | Alzheimer's disease; pathologic stage unknown | | | | ChP, FC, MesFat, SubFat |
| A.15 | 75 | F | 24h | (-) | Brochopulmonary hemorrhage | | | | ChP, FC, MesFat, SubFat, CSF |
| YA.10 | 73 | M | 4h | AD | | | | | ChP, FC |
| A.16 | 66 | F | 22h | (-) | Achondroplasia | | | | ChP, FC, MesFat, SubFat |

**Table S1:** Clinical and experimental metadata for donor samples used in this study. Abbreviations: **ChP**: choroid plexus; **AD**: Alzheimer's disease; **ADNC**: Alzheimer's disease neuropathologic change; LBD: Lewy body disease; **FC**: frontal cortex; **MesFat**: mesenteric fat; **SubFat**: subcutaneous fat; **CSF**: cerebrospinal fluid

| Cell type | High-expression marker genes |
|---|---|
| Endothelial | FLT1, PECAM1 |
| Glial | GFAP, SLC1A3, FABP7, BLBP |
| Ependymal | FOXJ1, CIMAP3, DYNLRB2 |
| Epithelial | TTR, KCNE2, KCNQ1, KCNA3, MUC1, EPCAM, KRT8, KRT18, AQP1, OTX2, CLDN1, CLDN2, CLDN3, SLC4A5, ATP1A1, PAX6, FOXJ1, LAMP5, PLTP, GPR125 |
| Macrophage | C1QA, C1QB, C1QC, CCR5, CD14, CD68, CD163, CD86, CLEC7A, CTSB, CX3CR1, DOCK2, DOCK4, FCGR1A, FCGR3A, ITGAX, MRC1, MSR1, PTPRC, TBXAS1, SLC8A1, TLR2 |
| Neural | NEFL, MAPT, KCNQ5, RBFOX3, MAP2, TUBB3, DCX, NEFM, SYN1, SLC17A7, GAD1, GAD2, STMN2, SNAP25, CAMK2A, ELAVL4, GRIN1 |
| Mesenchymal | NT5E, ENG, PDGFRA, COL1A1, COL1A2, DCN, LUM, ACTA2, TAGLN, RGS5, NOTCH3, MCAM, VIM, FN1, PDGFRB, THY1 |
| T-cell | CD3D, CD3E, CD3G, THEMIS, CAMK4, CD52, IL32, LTB, IL7R, CD2, INPP4B, TRAC, TC2N, ADAM19, TRAT1, SPOCK2, ICOS, CXCR4, CD27, ITM2A, CD247 |
| B-cell | MS4A1, CD19, EBF1, IGLV6-57, IGLV4-60, IGKV3D-20, IGKV6-21, IGKJ3, IGKJ4, KLHL14, CEP295NL, WNT16, GNG3, ICOSLG, OSBPL10, FCRL5 |
| Dendritic cell | CD1A, CLEC4C, NRP1, CR2, FCER2, IL3RA, CD207, ITGAX, HLA-DRA, HLA-DRB1, HLA-DRB3 |

**Table S2:** Cell type gating markers used for annotation of choroid plexus cell populations

| Module | Genes |
|---|---|
| Myeloid Identity | LYZ, CSF1R, SPI1, CD68, MPEG1 |
| Granulocytic | MPO, ELANE, S100A8, S100A9, FCGR3B |
| Monocytic lineage | CD14, FCGR3A, CCR2, CX3CR1 |
| Immature / progenitor | MKI67, TOP2A, LY6E, MPO |
| Circulating monocyte | S100A8, S100A9, CCR2, FCN1, VCAN |
| Tissue macrophage | MRC1, CD163, MSR1, MARCO, TREM2, APOE, FABP5, SPP1 |
| Inflammatory | IL1B, TNF, NFKBIA, CXCL9, CXCL10, STAT1, IRF1, GBP1 |
| Anti-inflammatory | MRC1, CD163, MSR1, C1QA, C1QB, C1QC, TREM2, APOE |
| Antigen presentation | HLA-DRA, HLA-DRB1, CD74, B2M, CIITA |
| Phagocytosis | MERTK, CD36, MARCO, MSR1, LPL, GPNMB |
| Brain microglia | TREM2, TMEM119, P2RY12, CX3CR1 |
| Liver Kupffer | CLEC4F, VSIG4, CD5L, MARCO |
| Lung alveolar | SIGLEC1, MARCO, FABP4, PPARG, LPL |
| Retina Choroid | CD163, MRC1, TREM2, APOE, SPP1, LYVE1, MERTK |

**Table S3:** Marker gene sets used to annotate myeloid lineage states, activation programs, and tissue-specialized macrophage populations.

| Module | Genes |
|---|---|
| TTN regulatory factors | BACH1, FOS, JUN, JUNB, HSF1, MEF2A, MEF2C, MEF2D, IRF8, MAFB, TCF12, PKNOX1 |
| Tissue Residence | CX3CR1, ITGAM, ITGAX, CSF1R, CD68, CD163, NR4A1, NR4A2, S1PR1 |
| DAM-like | TREM2, TYROBP, APOE, APOC1, LPL, GPNMB, SPP1, CST7, CTSB, CTSD, CTSS, LILRB4, AXL, CSF1R |
| Autophagy | ULK1, ULK2, ATG13, RB1CC1, ATG101, BECN1, PIK3C3, PIK3R4, ATG14, AMBRA1, WIPI1, WIPI2, ATG3, ATG4A, ATG4B, ATG4C, ATG4D, ATG5, ATG7, ATG10, ATG12, ATG16L1, GABARAP, GABARAPL1, GABARAPL2, MAP1LC3A, MAP1LC3B, SQSTM1, OPTN, TAX1BP1, NBR1, CALCOCO2, TBK1, IRGM, BNIP3, BNIP3L, VPS34, VPS15, UVRAG, RUBCN, ATG9A, ATG2A, ATG2B, TECPR1, EPG5, VPS11, VPS16, VPS18, VPS33A, VPS39, VPS41, RAB7A, RAB11A, RAB5A, RAB5B, RAB5C, RAB7B, SNAP29, STX17, VAMP7, VAMP8, LAMP1, LAMP2, CTSB, CTSD, CTSL, VPS4A, VPS4B, ATG9B, EPN1, DNM2, RUBCNL, UVRAG, KIAA1324 |
| Senescence | TP53, TP73, CDKN1A, CDKN1B, CDKN2A, CDKN2B, CDKN2C, RB1, RB2, E2F1, E2F2, E2F3, CCND1, CCND2, CCND3, CCNE1, CCNE2, CCNA2, CCNB1, CCNB2, CDK1, CDK2, CDK4, CDK6, MDM2, MDM4, GADD45A, GADD45B, GADD45G, PML, CDKN3, CHEK1, CHEK2, ATM, ATR, H2AFX, RAD51, RAD52, RAD9A, PCNA, LMNB1, LMNB2, HMGB1, HMGB2, IL6, IL8, CCL2, CCL5, CXCL1, CXCL2, CXCL3, CXCL5, CXCL8, TNF, IL1A, IL1B, MMP1, MMP3, MMP9, SERPINE1, IGFBP3, IGFBP7, CXCL12, CXCL16, CXCL10, CXCL11, IFNB1, IFNG, TGFBR1, TGFBR2, SMAD2, SMAD3, SMAD4, SMAD7, FOXO1, FOXO3, SIRT1, SIRT6, NAMPT, PARP1, PARP2, CXCR2, CXCR4 |
| Cytoskeleton | ACTB, ACTG1, TUBB, TUBA1A, TUBB2A, TUBB3, VIM, EZR, MSN, RDX, CFL1, CFL2, PFN1, WAS, WASL, ARP2, ARP3, ARPC1A, ARPC1B, ARPC2, ARPC3, ARPC4, ARPC5, DIAPH1, DIAPH2, FMNL1, FMNL2, FMNL3, MYH9, MYH10, MYH14, MYL6, MYL9, TPM1, TPM2, TPM3, TPM4, ADD1, ADD2, ADD3, CORO1A, CORO1B, CORO1C, PLEK, PLEKHA1, PLEKHA2, PLEKHB2, ITGA4, ITGAM, ITGAX, ITGB1, ITGB2, ITGB7, TLN1, TLN2, VCL, PXN, ZYX, RAC1, RAC2, CDC42, RHOA, RHOC, RHOG, DOCK2, DOCK8, ABI1, ABI2, NCK1, NCK2, CRK, CRKL, SOS1, SOS2, RACGAP1, ROCK1, ROCK2, LIMK1, LIMK2, CFL1, CFL2, DSTN, PFN1, PFN2, MYO1E, MYO9B, MYO10, MYO1F, MYO7A, MYO18A, MAP1A, MAP1B, MAP1S, MAP2, MAP4, MAP6, MAP7, MAPK1, MAPK3, MAPK14, MAPK8, MAPK9, MAPK10, PAK1, PAK2, PAK3, PAK4, PAK6, PAK7, KIF1B, KIF5B, KIF11, KIF23, DYNLT1, DYNC1H1, DCTN1, DCTN2, DCTN3, DCTN4, DCTN5, DCTN6, FLNA, FLNB, FLNC, SPTAN1, SPTBN1, SPTBN2, SPTBN4, DBN1, DBN2, FHOD1, FHOD3, LCP1, LCP2 |
| Oxidative Phosphorylation | NDUFA1, NDUFA2, NDUFA3, NDUFA4, NDUFA5, NDUFA6, NDUFA7, NDUFA8, NDUFA9, NDUFA10, NDUFA11, NDUFB1, NDUFB2, NDUFB3, NDUFB4, NDUFB5, NDUFB6, NDUFB7, NDUFB8, NDUFB9, NDUFB10, NDUFB11, NDUFC1, NDUFC2, NDUFS1, NDUFS2, NDUFS3, NDUFS4, NDUFS5, NDUFS6, NDUFS7, NDUFS8, SDHA, SDHB, SDHC, SDHD, UQCRC1, UQCRC2, UQCRFS1, CYC1, COX4I1, COX5A, COX5B, COX6A1, COX6B1, COX6C, COX7A2, COX7B, COX7C, ATP5F1A, ATP5F1B, ATP5F1C, ATP5F1D, ATP5F1E, ATP5MC1, ATP5MC2, ATP5MC3, ATP5ME, ATP5MF, ATP5MG, ATP5PB, ATP5PD, ATP5PF, ATP5PO, ATP5D, ATP5J, ATP5J2, ATP5L, ATP5O |
| Phagocytosis | FCGR1A, FCGR1B, FCGR1C, FCGR2A, FCGR2B, FCGR3A, FCGR3B, CD36, MSR1, MRC1, ITGAM, ITGB2, LYN, SYK, HCK, LCP2, VAV1, RAC1, RAC2 |
| MHC class II | HLA-DRA, HLA-DRB1, HLA-DRB5, HLA-DPA1, HLA-DPB1, HLA-DQA1, HLA-DQB1, CIITA, CD74, CD86, CD80, CD40, TAP1, TAP2, B2M, LCP2, LY96, IRF1, IRF8, STAT1 |
| Reactive Oxygen Species | SOD1, SOD2, CAT, GPX1, GPX3, GPX4, PRDX1, PRDX2, PRDX3, PRDX4, PRDX5, PRDX6, TXN, TXN2, TXNRD1, TXNRD2, NQO1, HMOX1, GSR, GCLC, GCLM, GSS, GLRX, GLRX2, SRXN1, SESN1, SESN2, KEAP1, NFE2L2 |
| Efferocytosis | MERTK, AXL, TYRO3, GAS6, PROS1, LRP1, CD36, ITGAV, ITGB3, MFGE8, TIMD4, STAB1, STAB2 |

**Table S4:** Macrophage functional gene modules. Abbreviations: DAM = disease associated microglia; MHC = major histocompatilbility complex.

### 2 Supplementary Methods

#### 2.1 Key reagents

| Reagent/Resource | Source | Identifier | |
|---|---|---|---|
| PCR primers | Integrated DNA Technologies Coralville, IA | 5'-ATGAGCAATGGAGACCTAACAG-3' | (Primer 1, forward) |
| | | 5'-CCGAAGCCAAAGTCAAGATCC-3' | (Primer 1, reverse) |
| | | 5'-CAGGTTCCTTCTTAGGCACAG-3' | (Primer 3, forward) |
| | | 5'-GAGGTGTCAGTCACAGTCCC-3' | (Primer 3, reverse) |
| | | 5'-GCAGCCTCTTCCTTAGGTGC-3' | (Primer 7, forward) |
| | | 5'-CAAACCACCTCCTGTGGAACC-3' | (Primer 7, reverse) |
| | | 5'-CTGCTGCGACACTCTATGAC-3' | (Primer 12, forward) |
| | | 5'-CTAAATTAACCATCCGTGAAACC-3' | (Primer 12, reverse) |

**Table S5:** Key resources table

### 3 Supplementary Figures

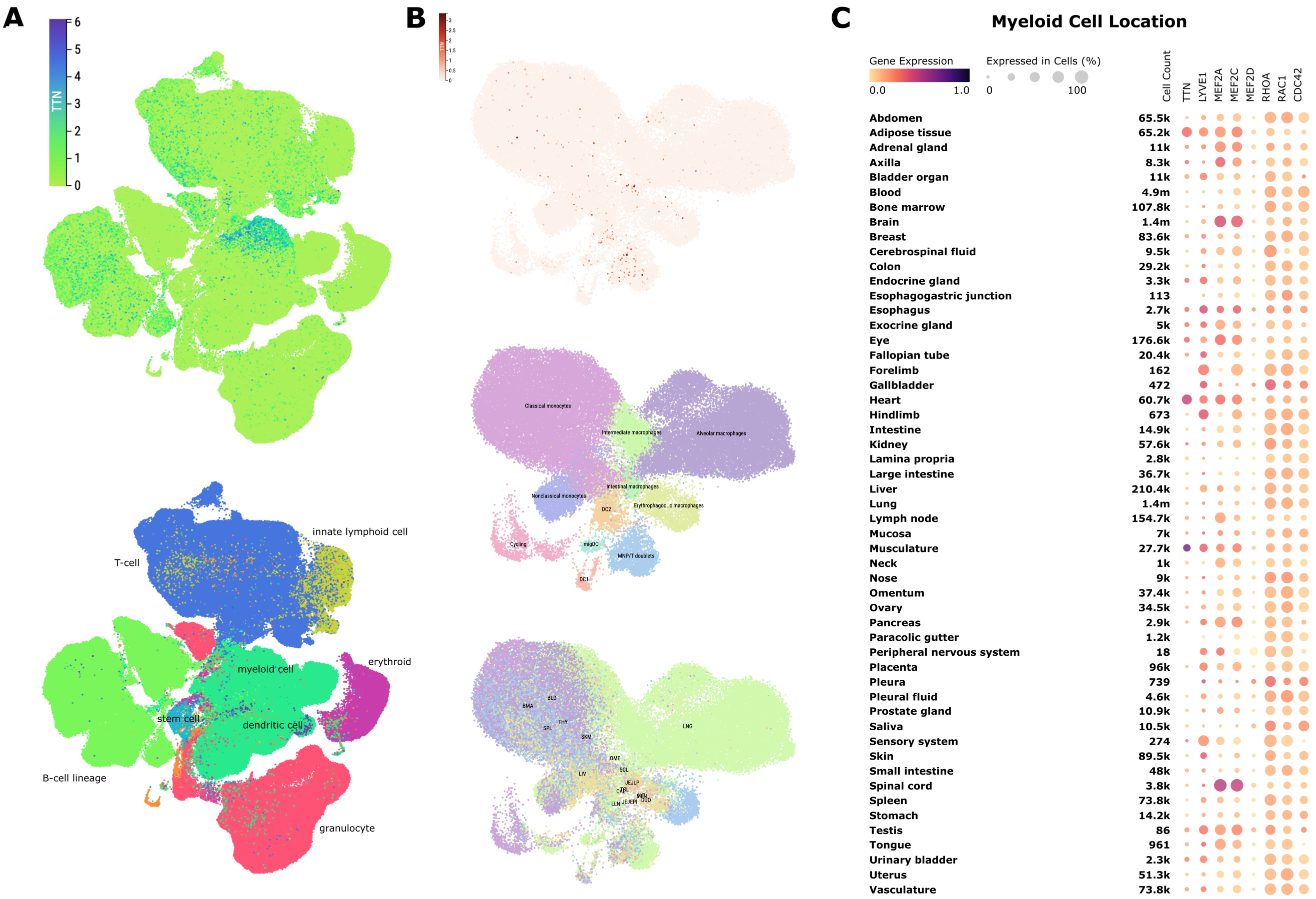


**Figure S1: TTN-expression in macrophages in other body tissues**. **(A)** Single cell data from the immune cell collection of the Tabula Sapiens [1] reference human cell atlas v2.0 [2], projected onto UMAP coordinates and labeled by TTN expression (top) and immune cell type (bottom). Expression is observed predominantly within the myeloid cell cluster, along with lymphoid subpopulations. **(B)** Single cell data from the cross tissue immune cell atlas [3], labeled by TTN expression (top), immune cell subtype (middle), and organ of residence (bottom). In addition to myeloid lymphocyte doublets, TTN epxression is seen in intestinal macrophages. Abbreviations: BLD=blood, BMA=bone marrow, SPL=spleen, THY=thymus, SKM=skeletal muscle, LIV=liver, OME=omentum, LNG=lung, SCL=sigmoid colon, CAE=caecum, JEJLP=jejunum lamina propria, JEJEPI = jejunum epithlium, LLN=lung draining lymph nodes, MLN = mesenteric lymph nodes. **(C)** Expression of TTN and other mechanosensitive genes in myeloid cells by anatomic location, in [2]. In addition to those occupying skeletal muscle and heart, TTN expression in myeloid cells is elevated in adipose tissue

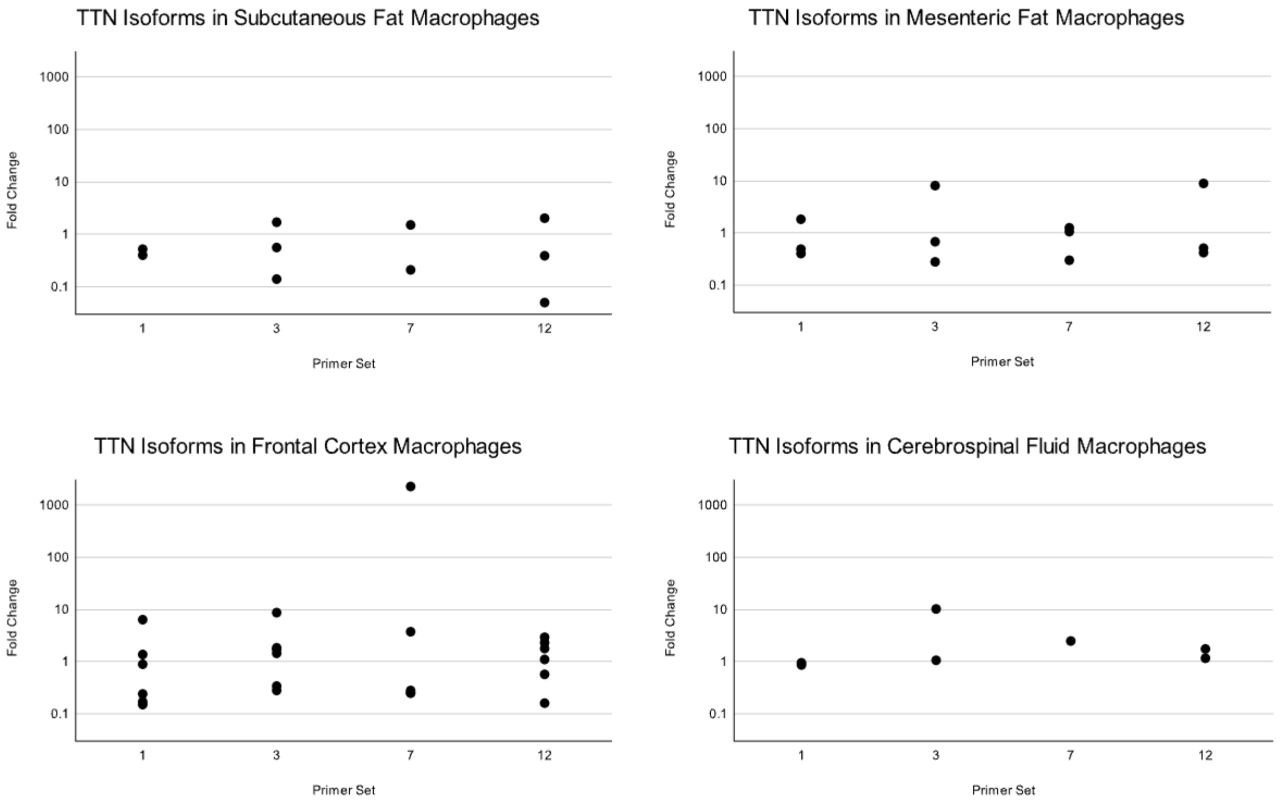


**Figure S2:** qPCR on all tissues. Macrophages were selected for using CD163 MicroBeads (see Methods). TTN primers 1, 3, 7, and 12 were used to amplify gene expression in human subcutaneous fat, mesenteric fat, frontal cortex, and CSF macrophages. Data points represent fold changes, computed from $\Delta\Delta$Cq, in TTN isoform expression, normalized to GAPDH, in CD163-bead enriched macrophages relative to whole tissue. The x-axis represents different primer sets (1, 3, 7, and 12), and the y-axis shows fold changes (log scale). Each point represents a single donor sample/primer set pair. TTN transcript enrichment is noted in macrophages isolated from frontal cortex, CSF, and, to a lesser degree, mesenteric fat.

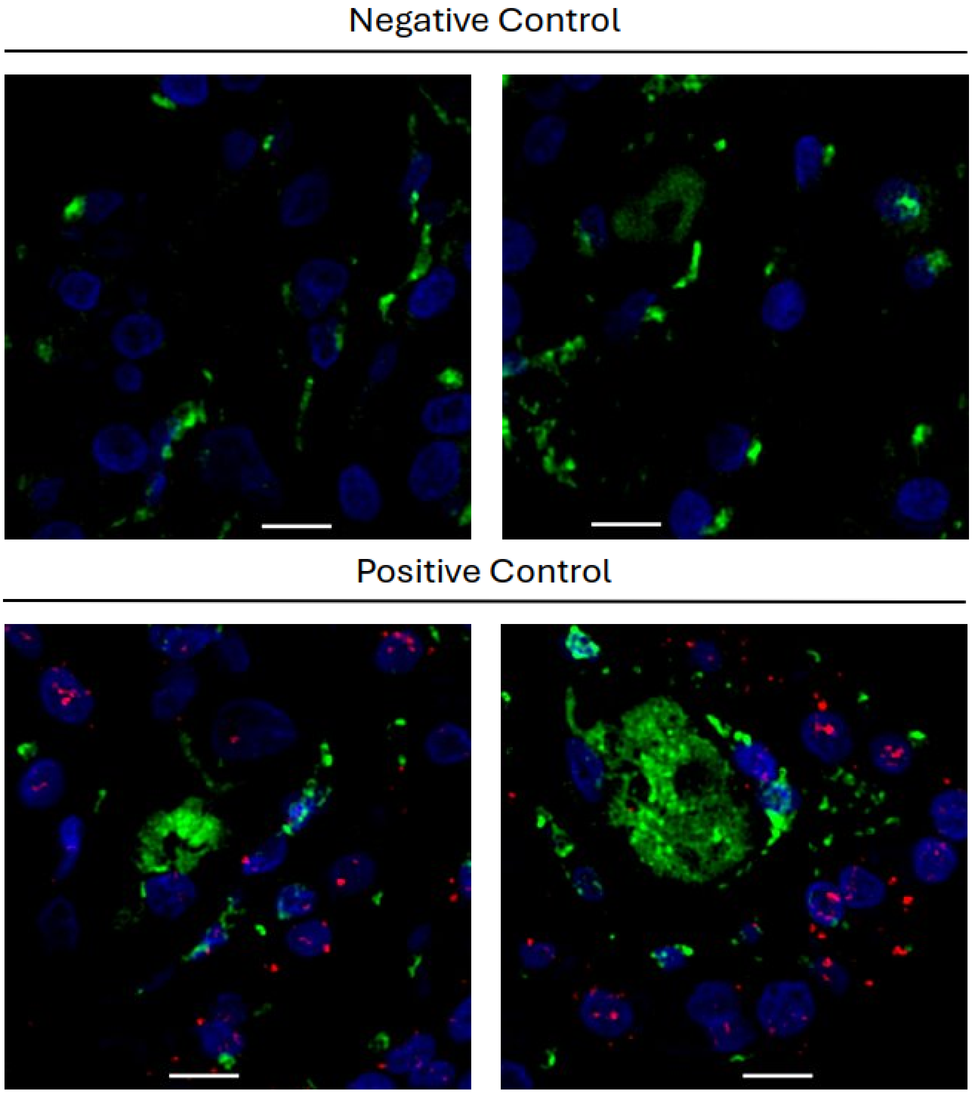


**Figure S3:** RNAscope Controls on human choroid plexus FFPE tissue. Negative control tissues were incubated with RNAscope 3-plex Negative Control Probe (ACDBio, #320871) for dapB. Positive control received RNAscope 3-plex Positive Control Probe (ACD Bio, #320861) with probes for C1-POL2A. Red = RNAscope probe. Green = Immunofluorescence with Iba1 primary antibody. Scale bars = 10 µm

### References (Supplementary Information)


1. THE TABULA SAPIENS CONSORTIUM. The Tabula Sapiens: A multiple-organ, single-cell transcriptomic atlas of humans. *Science* **376.** Publisher: American Association for the Advancement of Science, eabl4896. https://www.science.org/doi/10.1126/science.abl4896 (2025) (May 2022).

2. Consortium, T. T. S. & Quake, S. R. *Tabula Sapiens reveals transcription factor expression, senescence effects, and sex-specific features in cell types from 28 human organs and tissues* en. ISSN: 2692-8205 Pages: 2024.12.03.626516 Section: New Results. Aug. 2025. https://www.biorxiv.org/content/10.1101/2024.12.03.626516v2 (2025).

3. Domínguez Conde, C. *et al.* Cross-tissue immune cell analysis reveals tissue-specific features in humans. *Science* **376.** Publisher: American Association for the Advancement of Science, eabl5197. https://www.science.org/doi/10.1126/science.abl5197 (2025) (May 2022).
